# Emergent control of ant learning walks using the mismatch between path-integration and visual cues

**DOI:** 10.64898/2026.05.29.728637

**Authors:** Thomas Misiek, Andrew Philippides, Paul Graham, James Knight

**Affiliations:** Sussex AI, School of Engineering and Informatics, University of Sussex, Brighton, UK; School of Life Sciences, University of Sussex, Brighton, UK

## Abstract

Despite their small brains, desert ants can safely navigate back to their nest after travelling hundreds of metres to find food. This ability relies on visual memories, gathered on initial *Learning Walks* (LWs), during which ants slowly explore the surroundings of the nest. LWs follow a clear progression: early walks are short, spiraling, and interspersed with pirouettes: brief stops during which the ant perform a visual scan. Successive LWs become straighter and extend farther from the nest, until they transition into foraging trips. However, when the visual panorama changes, LWs reappear. Existing models explain *how* stored panoramic views can guide homing, but not *when* visual memories are collected or how LW dynamics are controlled. Here, we propose that a single error signal – generated from the mismatch between the estimated nest direction provided by path integration and visual predictions – scaffolds visual learning and drives LW dynamics. Using 3D reconstructions of desert ant experiments, we show that this mechanism reproduces the main features of LWs, including the transition to foraging and reoccurrence of LWs after environmental change. Our model is biologically plausible and consistent with known insect navigation circuits.

**Author summary:** A tiny nest entrance in the desert is easy to miss, but doing so may have lethal consequences for a foraging desert ant, returning home after a long scavenging trip. To solve this problem, desert ants spend several days performing Learning walks (LWs) to learn the visual landmarks around their nest before they begin searching for food. LWs are structured active learning procedures and clear patterns can be identified across ant species. LWs take the form of nest-centred spirals, interspersed with frequent visual scans. Over two to four days, LWs become longer, straighter and walking speed increases, before the ant finally transitions to foraging. LWs may reoccur even in experienced foragers, when a new landmark is introduced near the nest.

No previous model has provided a mechanism for how LWs are controlled. Here, we propose that the transition from LWs to foraging is driven by visual familiarity. In our model, unfamiliar surroundings promote learning and exploration; as the visual panorama becomes familiar, behaviour shifts to foraging. Using 3D simulations of experimental layouts, we demonstrate that LWs can emerge from simple rules, and that our simulated agents reproduce the main features of LWs observed in desert ants.

## Introduction

Central-place foraging insects spend much of their lives efficiently navigating between food sources and their nest [1–3]. Desert ants, for example, travel hundreds of meters from their nest to scavenge for dead insects and can reliably return home, despite limited neural resources [4–6]. Rather than relying on a single mechanism, ants combine complementary navigation strategies that operate redundantly and over different spatial scales [7–10].

In desert ants, an innate Path Integration (PI) system, driven by celestial [11], magnetic [12] and speed information [13, 14], continuously estimates a nestward vector, enabling homing even in unfamiliar terrain [13, 15–18]. However, PI progressively accumulates error as the outbound path lengthens, leading to growing uncertainty in the nest vector [17, 19]. Consequently, PI alone cannot reliably pinpoint the nest entrance after extended foraging excursions. Upon reaching the nest position estimated by PI, ants switch to systematic search: walking in expanding loops centred on the predicted nest location. [17, 20–24]. However, these searches are slow and energetically costly, increasing exposure to heat, desiccation, and predation. Therefore, in the vicinity of the nest, Ants supplement PI with a variety of mechanisms to reduce search time. These include innate responses such as following downwind CO_2_ plumes from the nest [10], as well as learned strategies like memorising local olfactory cues [25–28], and exploiting vibration, magnetic, and tactile cues when available [29, 30].

Desert ants also allocate significant neural resources to visual learning. This is reflected in their well-developed compound eyes and optic-lobes [31, 32], and mushroom bodies [33]. These adaptations help desert ants to find their nest holes [3, 25, 34], with some species even constructing nest hills to improve the visibility of the nest in environments lacking surrounding terrestrial cues [35]. *Snapshot navigation* is a hypothesis as to how the visual information acquired by insects is used for homing. It relies on the fact that the visual discrepancy between two panoramic views increases smoothly with both translational and rotational displacement [36]. Therefore, navigation can proceed by minimizing visual mismatch between the current view and stored views, allowing the original acquisition position and orientation to be recovered [37, 38]. Visual Compass (VC) models implement this hypothesis by having agents rotate on the spot and compare their current view against stored nest-oriented snapshots. This process generates a Rotational Image Difference Function (RIDF), the minimum of which predicts the nest azimuth and supports visual homing [36, 39, 40]. The same principle can enable route navigation by aligning the animal with a familiar heading sequence [39, 41]. VC navigation can be implemented algorithmically by directly computing similarity between images using a function such as Mean Squared Error (MSE) [36] or by using an Artificial Neural Network [39, 42–44]. In ants, the visual memories supporting the VC could be stored in the Mushroom Body (MB) – a key centre for multimodal learning and memory [45–49]. On this basis, several VC implementations have been developed using models of the Mushroom Body [50–52].

In order to acquire the visual information about the environment surrounding the nest required for homing – including local landmarks and the skyline – ants perform structured exploratory excursions known as Learning Walks (LWs) [4, 53–57]. Ants invest substantial effort in LWs: they perform two to seven LWs, across two to four days before starting foraging [54, 58]. During LWs, ants do not collect food but instead slowly sample the panorama, acquiring views that will later support homing [53, 54]. While it has been shown that ants’ visual homing performance improves with experience and successive LWs [53, 54], it is still unclear when during the LWs ants capture the visual memories that will later enable visual homing. In desert ants LWs are characterised by continuous almost 180*^◦^* oscillations [54], as well as frequent discrete scans called pirouettes, and rarer, small walked circles called voltes [59]. Pirouettes are tight rotations about the vertical body axis, observed in many desert ants, during which the ant stop moving forward, turn around to face the nest and then back toward the initial direction to resume walking. Pirouettes also usually contain multiple shorter stopping phases (≥ 100 ms) in different directions [59]. Early studies proposed that hymenopterans memorise views primarily when facing the nest. A clear bias toward fixating the nest was first observed in flying insects during learning flights [38, 60–62], and was one of the inspirations for snapshot navigation [36]. However, in desert ants, while there is some evidence that the *longest* fixation during pirouettes is usually oriented toward the nest [55, 59], pirouette contains 4 ± 3 fixations [59] in multiple directions. Müller and Wehner [55] hypothesized that nest fixations during pirouettes are guided by the PI signal. They showed that in the Namibian desert, *Ocymyrmex robustior* can turn toward the nest direction even when the entrance is not visible, suggesting that PI provides a nestward directional signal that orients pirouettes during LWs.

The geometry of LWs varies widely between species, ants of the same colony and across successive trips by the same ants [54, 58, 63]. Yet, despite this variability, LWs share a robust set of characteristic features across desert ants species [54]:

### Increasing walking speed

Ants average walking speed is slower during early LWs compared to later LWs and foraging [58, 64, 65].

### Decreasing sampling and pirouettes

Early LWs contain frequent nest-directed pirouettes. [59]. In subsequent LWs, the total number of pirouettes per walk decreases [55] as well as their frequency [66].

### Straightening walks

Ants tend to spiral around the nest during early LWs but later LWs become straighter [53, 55].

### Expanding walks

On average, successive LWs increase in duration, extend farther from the nest, and cover an increasingly larger area [53, 63].

### Homing performances improve with experience

Experienced zero-vector ants show higher search accuracy, centring their searches closer to the nest than näıve zero-vector ants during homing tests [53].

### Bringing back food

Experienced ants are more likely than näıve ants to pick-up and bring back food they find [53].

### First foraging walk and transition to exploitation

Ants start foraging after between two to seven LWs [53, 58, 66] with substantial variability between ants from the same nest.

### Reappearance of learning after change

When the visual panorama is altered, by adding a cylinder or removing trees close to the nest for example, learning behaviours like pirouettes or even full LWs reoccur immediately [55, 65, 67–69].

Together, these dynamics lead LWs to transition into mature foraging trips: straight, efficient, and food oriented. While the snapshot navigation hypothesis explains how stored nest-directed views can support visual homing, it remains unclear how visual memories are dynamically acquired during LWs. In particular, what triggers pirouettes? What mechanisms govern the progressive expansion of LWs, and what controls the transition from spiralling to straighter walks? Finally, what explains the reoccurrence of LWs following environmental change? One possibility is that these transitions are governed by an intrinsic schedule – such as a preset number of LWs to perform, and planned expansion of the LWs. Alternatively, the transition from foraging to LWs could be governed by a simple environmental signal such as cumulative light exposure.

Repeated light exposure for a period of two to four days is known to induce MB synaptic reorganization [70, 71], during the early foraging careers of ants so would be a candidate mechanism for triggering other physiological and behavioural changes. However, altering the landscape, by adding landmarks or felling trees around the nest makes learning behaviours reoccur immediately in experienced ants [55, 65, 67–69], which is difficult to reconcile with a simple intrinsic schedule, or a cumulative environmental signal. We hypothesize that LW expression is regulated online by an internal error signal that continuously evaluates the reliability of the Visual Compass and triggers learning when VC predictions become unreliable. One possible candidate for such an error signal would be to the ‘strength’ of the familiarity signal provided by the VC. Previous work has used such signals to weigh the VC signal in multimodal navigation [72] or to detect changes in the visual panorama and modulate exploratory behaviours [68]. Yet, visual familiarity alone is ill-suited to assess the accuracy of directional predictions. Indeed, a strong familiarity signal may not be associated with a correct directional estimate: two views sampled in close proximity, but acquired on opposite sides of the nest may look similar but should generate opposite steering predictions. We instead propose that the angular discrepancy between the PI home vector and the VC predictions provides a more reliable error signal during LWs. We call this angular error *E*_PI_*_/_*_VC_ throughout the paper (Fig. 1). Averaged over time, this signal provides a measure of the familiarity of the visual environment and the reliability of the VC.

**Fig 1.**
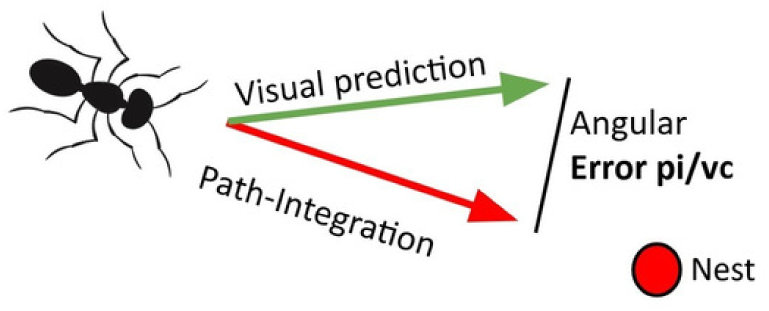
**Visualization of the angular error (***E*_PI_*_/_*_VC_). In our model, all learning behaviours of LWs are modulated by the mismatch between PI and visual cues: *E*_PI_*_/_*_VC_. High error triggers pirouettes and spiralling behaviour. PI as a red arrow. VC prediction as a green arrow. Nest in red.

In our model, the error signal *E*_PI_*_/_*_VC_ regulates key features of LWs: higher mismatch increases the probability of triggering a pirouette, decreases walking speed, promotes spiralling behaviour, and can trigger premature return to the nest. Using simulations of classic desert ant experiments [53, 55, 65], we reproduce the eight LW properties described above and demonstrate that robust and adaptive control of LWs can come from a single biologically plausible error signal that is compatible with known neuroanatomy of insect navigation circuits.

## Results

### The active learning model

In this section, we will show that the angular error signal *E*_PI_*_/_*_VC_ can modulate multiple sensorimotor motifs of learning and reproduce the main empirical properties of LWs listed above. We first briefly describe the Visual Compass and PI models that we use in all simulated experiments, and then describe how they interact through the active learning model. Full model specifications and implementation details are provided in the Methods section.

### Visual-Compass model

Fleischmann et al. [59] found that, although each pirouettes made by (*Cataglyphis noda*) ants include prolonged nest-facing stopping phases, they also make 4 ± 3 fixations across a broad azimuthal range. This observation suggests that snapshot acquisition might not be restricted to nest-facing views and that visual information across multiple orientations could be acquired during learning. Based on these observations, we implement our Visual Compass using a model of the Mushroom Body which captures panoramic snapshots across a distribution of azimuths (Fig. 2B-C and see Methods for full model description). This model follows a similar principle to the model presented by Wystrach et al. [51, 73], in which visual memories were distributed between two distinct MBONs representing whether the goal lies to the left or right of the agent’s current heading. However, in our model, views are encoded at a finer angular resolution using eight MBONs, each associated with a preferred egocentric azimuth of the nest (Fig. 2G). The MBONs form a circular arrangement with an angular spacing of 45*^◦^* to match the columnar layout of the CX. Similarly to previous MB models, learning is implemented via an anti-Hebbian rule at KC–MBON synapses (Fig. 2D-E-F) [50, 51]. Similarly to Wystrach et al. [51, 73], the PI signal gates learning to a single MBON at a time: in our model, this is the MBON associated with the egocentric nest direction signalled by the corresponding CX column

**Fig 2.**
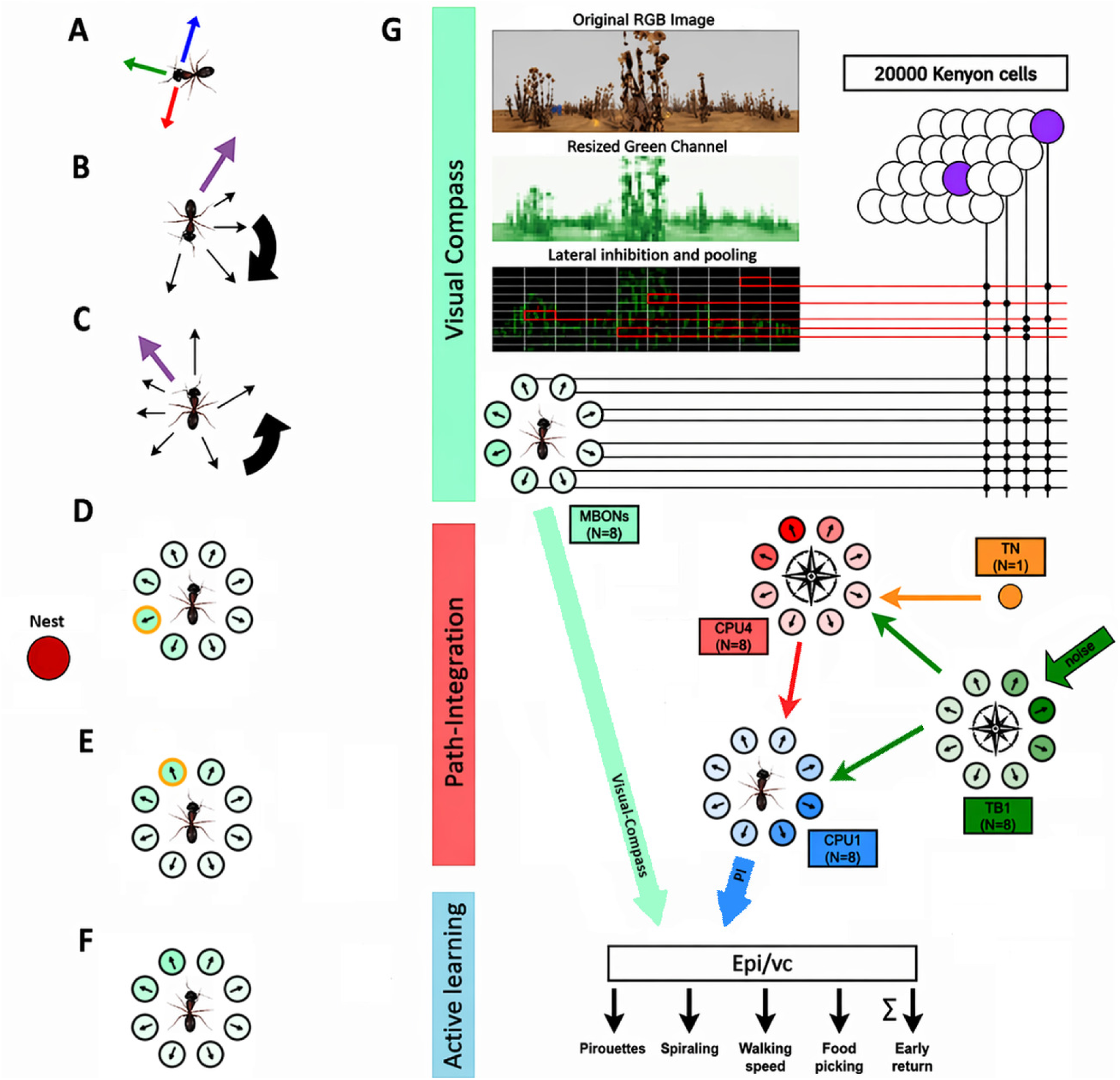
The active learning model. In A-G, the nest is the red circle near D. **A** Red: home vector. Green: spiraling vector. Blue: anti-home vector. **B** Partial pirouette. Purple arrow: walking direction of the agent before the pirouette is initiated. Thin black arrows: snapshots directions. Large black arrow: rotational direction of the pirouette. **C** Representation of a full pirouette with 6 fixations. **D** During learning (i.e., fixation during a pirouette), PI controls plasticity. Only the KC-MBON weights of the MBON associated with the current egocentric nest-orientation are updated. Turquoise: MBONs. Orange: selected MBON. **E** Another example, with a different egocentric direction of the nest. **F** During inference, the visual compass prediction corresponds to the direction of the MBON with the lowest activation (greener = lower activation in this illustration). **G** Diagram of the model. **Top:** Visual processing of the image and visual compass. The original RGB image is downsampled, and the green channel is isolated. Lateral inhibition is computed using a center-surround mechanism, and the low dimensional VPN output vector is created by pooling over a 10 × 10 grid on the lateral inhibition array. The 100 VPNs are sparsely connected to a population of 20,000 KCs (purple); each KC forms five random synapses with the VPNs. The KC population is sparsely activated, with 5% of KCs active at any time. The population of KCs is fully connected to the eight directionally tuned MBONs (turquoise). **Middle:** The TB1 ring (compass, green) receives noisy input. The TB1 ring and TN1 neuron (encoding stride, orange) update the allocentric PI memory in the eight CPU4 neurons (red). Allocentric PI information and compass information are used to convert the PI signal into the egocentric frame in CPU1 (blue). **Bottom:** The egocentric PI signal and visual compass signal are used to compute *E*_PI_*_/_*_VC_. *E*_PI_*_/_*_VC_ probabilistically triggers pirouettes and spiralling behaviour. Walking speed is inversely proportional to *E*_PI_*_/_*_VC_. *E*_PI_*_/_*_VC_ is integrated in an accumulator and triggers homing when above a threshold, as well as food acceptance.

During inference, a winner-take-all mechanism enables the least active MBON in the ring to bias the CX heading estimate toward the direction it encodes. In effect, a RIDF is computed ‘in-silico’ across the eight parallel MBONs, using the multi-directional snapshots captured during LWs. On the one hand, this implementation provides predictions with a finer angular resolution than the lateralized architecture of Wystrach et al. [51, 73], on the other hand, it does not require the inference-time rotational scans of the classic VC model [39]. Essentially, this is because these scans were performed during the pirouettes of LWs. Eliminating physical scans in our model enables continuous and precise VC predictions, which are critical to update the *E*_PI_*_/_*_VC_ signal.

### Path-Integration model

We use a simplified version of the anatomically-constrained PI circuit developed by Stone et al. [74]. PI is noisy and accumulates errors, so we add noise to the compass signal to fit experimental levels of PI error [55, 59] (see Methods). In our model, PI serves four functions:

1. It provides an initial homing vector that allows return to the nest before visual memories are learned [16–18].
2. The homing vector can be inverted to produce outward motion during the initial phase of LWs and rotated by 90*^◦^* to generate spiraling behaviour.
3. PI contributes to computing the mismatch between visual predictions and the PI-derived goal direction.
4. During learning, PI gates KC-MBON learning, selectively reinforcing the MBON associated with the correct egocentric nest direction.

### Expression of sensorimotor motifs of learning

In our simulations, naive agents depart from their nest and perform a number of consecutive walks. At the onset of a walk, the agent exits the nest in a random initial direction. The default behaviour of the agent is to follow the anti-nest direction provided by its PI (e.g., blue arrow in Fig. 2A). However, the *E*_PI_*_/_*_VC_ mismatch modulate the agent behaviour in multiple ways:

#### Walking speed

Forward speed is inversely proportional to the PI-VC mismatch, with a residual speed of 0.1 and a maximum speed of 1.1 centimeter per timestep when *E*_PI*/*VC_ is 0.

#### Spiralling

LWs have been described as spirals, forming arcs around the nest, with the ant walking perpendicular to the nest entrance [54]. To replicate this behaviour, in our model, *E*_PI_*_/_*_VC_ controls the probability of success of a per-timestep Bernoulli trial. If a trial succeeds i.e. there is high mismatch, spiralling is promoted by temporarily setting the goal direction perpendicular to the homing vector (e.g., green arrow in Fig. 2A).

#### Pirouettes

Mismatch also controls the success probability of another per-timestep Bernoulli trial which determines whether a pirouette is triggered. During pirouettes, the agent temporarily stops walking, rotates on the spot and acquires snapshots in multiple orientations, with around 4 ± 3 snapshots captured per pirouette (Fig. 2B, C).

#### Early return

Once the cumulative mismatch *C*_PI_*_/_*_VC_ exceeds a threshold, the agent switches to following the homing vector (e.g., red arrow in Fig. 2A). This mechanism helps prevent the agent from leaving the nest vicinity during early LWs, when the Visual Compass is still unreliable and *E*_PI_*_/_*_VC_ accumulates quickly. It limits PI error buildup before visual memories can reliably supplement PI. *C*_PI_*_/_*_VC_ is reset at the end of a walk and visual memories are consolidated upon return to the nest.

### Simulation of behavioural experiments

In this section, we simulate three desert ant experimental protocols (Müller and Wehner’s [55], Fleischmann et al. [53] (Fig. 3A, B) and Vermehren et al. [65] (Fig. 10A)) using 24 agents and up to nine consecutive walks. We show that our model closely replicates the behavioural trends they reported. All experiments were simulated using the same model’s parameters. Only the simulated layout and the number of simulated walks differed.

**Fig 3.**
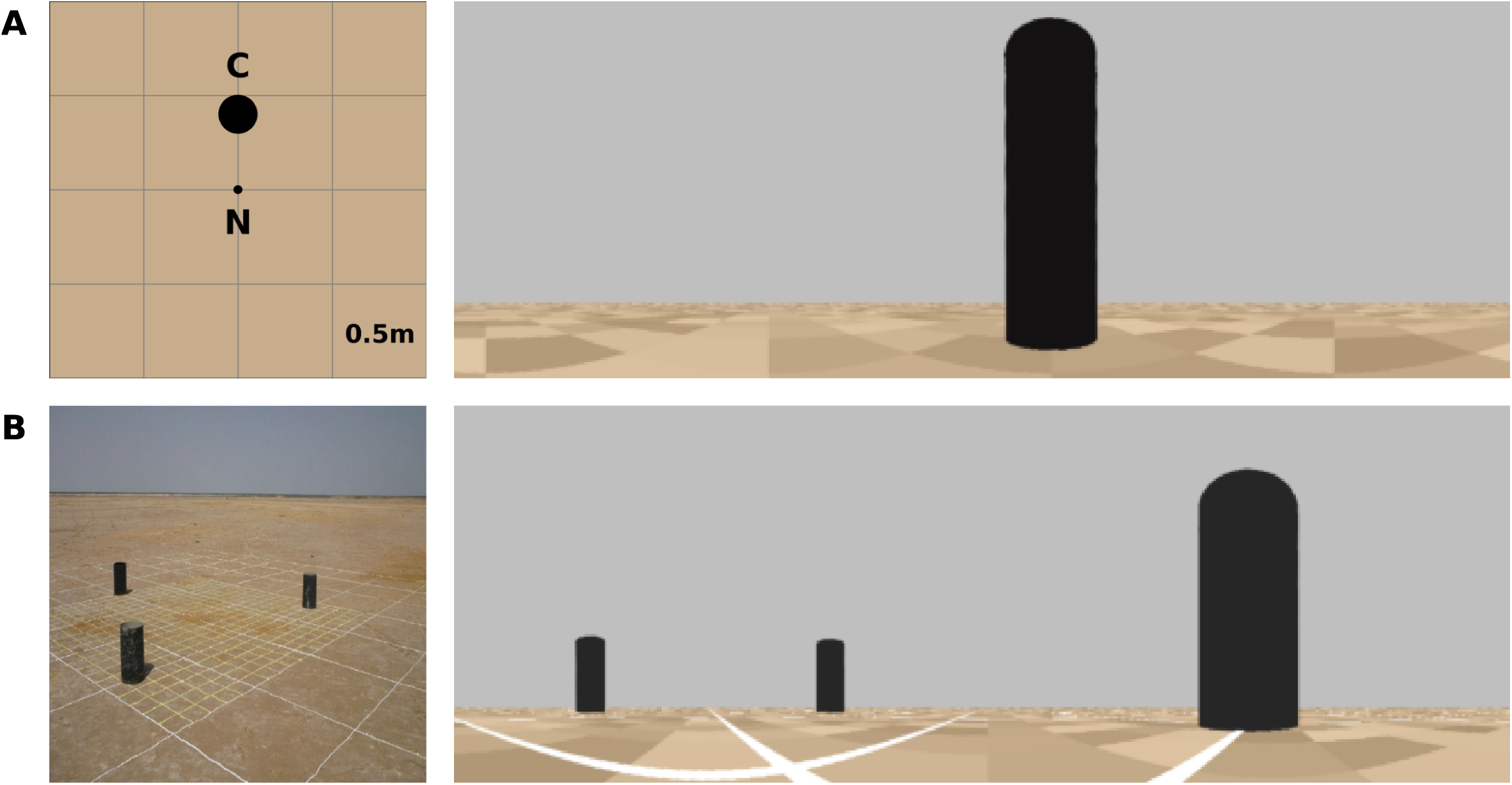
Experimental layouts. **A:** Müller and Wehner [55] layout (left) and corresponding 360*^◦^*panoramic render from our simulation (right). A single black cylindrical landmark (height 30 cm, diameter 11 cm) is positioned 40 cm north of the nest. View taken at the nest position, 5 cm above ground level. **B:** Fleischmann et al. [53] layout (left) and corresponding 360*^◦^* panoramic render from our simulation (right). Three black cylindrical landmarks (height 38 cm, diameter 22 cm) arranged in a triangular configuration, each located 2 m from the nest. View taken 1 m from the nest.

#### PI/VC mismatch decreases with successive LWs

Before presenting further results, we first verify that *E*_PI_*_/_*_VC_ decreases across successive walks, as this error signal modulates all sensorimotor motifs of learning in the model. From Fig. 4A, we observe that PI/VC error decreases across successive walks in both simulation layouts (Fig. 3). This is statistically supported by a one-way ANOVA (Müller: *F* = 15.06, *p* = 1.21 × 10*^−^*^13^; Fleischmann: *F* = 12.56, *p* = 1.36 × 10*^−^*^11^).

**Fig 4.**
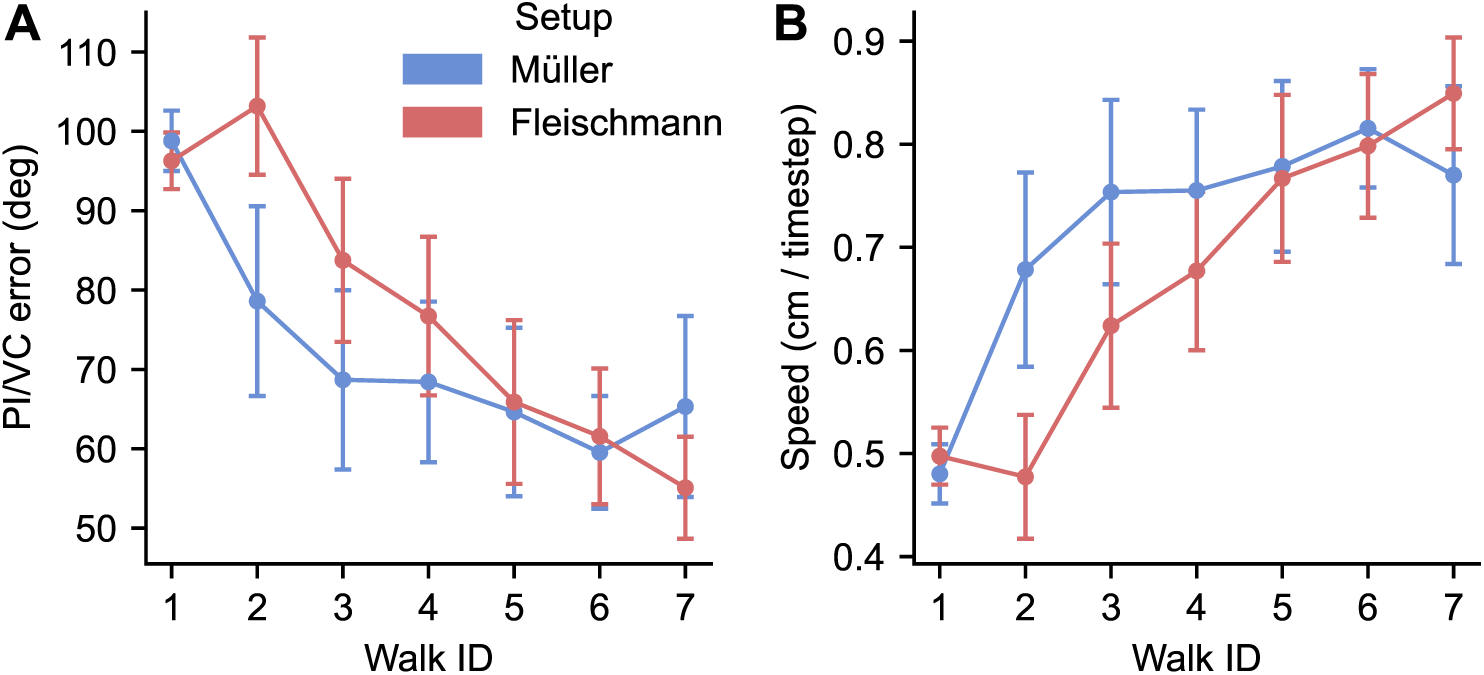
PI/VC mismatch decreases and walking speed increases across successive walks. **A:** PI/VC mismatch decreases across successive walks in simulations of both layouts (Müller in blue, Fleischmann in red). The error decreases faster in the Müller environment, but both converge to similar error levels after ∼five walks. The y-axis shows the absolute angular error between PI and VC predictions. **B:** Walking speed increases across successive walks in simulations of both layouts (Müller in blue, Fleischmann in red), with a steeper increase in the Müller environment, but both reach similar walking speed after ∼five walks. Speed is given in cm per timestep. Error bars indicate 95% confidence intervals of the mean.

Error decreases more rapidly during the early walks in the Müller environment than in the Fleischmann environment, before reaching a similar plateau in both conditions. Consistent with this, error differs significantly between conditions during the early phase (walks 1–4; two-sample *t*-test: *t* = −5.28, *p* = 2.70 × 10*^−^*^7^), but not during the late phase (two-sample *t*-test: *t* = −0.63, *p* = 0.534), indicating that agents in both environments converge to similar levels of performance at later stages of learning. We also find that *E*_PI_*_/_*_VC_ is higher during LWs than during foraging walks in both simulation layouts. In this paper, we classify walks using a distance threshold: those beyond 3 m from the nest are foraging walks, while those within 3 m are LWs. In the Müller environment, the mean error is significantly higher, with 99.58*^◦^* during LWs compared to 37.43*^◦^*during foraging walks (two-sample *t*-test: *t* = 17.83, *p* = 8.39 × 10*^−^*^25^). A similar pattern is observed in the Fleischmann environment, where the mean error is 84.06*^◦^* during LWs and 52.09*^◦^* during foraging walks (two-sample *t*-test: *t* = 13.05, *p* = 9.08 × 10*^−^*^27^).

This gradual decline in *E*_PI_*_/_*_VC_ provides the mechanism by which learning walks are suppressed with experience. It enables the transition to foraging and, in the following sections, we show how the decline in *E*_PI_*_/_*_VC_ allows the model to reproduce the key features of LWs.

#### Walking speed increases with successive walks

As walking speed is inversely proportional to *E*_PI_*_/_*_VC_, unsurprisingly, we observe that it increases across successive walks in both simulation layouts (Fig. 4B). This variation is statistically supported by a one-way ANOVA (Müller: *F* = 17.57, *p* = 1.35 × 10*^−^*^15^; Fleischmann: *F* = 13.82, *p* = 1.23 × 10*^−^*^12^).

Speed is also lower during LWs than during foraging walks in both simulation layouts. In the Müller environment, mean speed is 0.49 cm timestep*^−^*^1^ during LWs and 1.02 cm timestep*^−^*^1^ during foraging walks (*t* = −18.52, *p* = 1.71 × 10*^−^*^25^). Likewise, in the Fleischmann environment, mean speed is 0.64 cm timestep*^−^*^1^ during LWs and 0.86 cm timestep*^−^*^1^ during foraging (*t* = −13.63, *p* = 1.38 × 10*^−^*^28^; Fig. 4B).

Müller and Wehner [55] and Fleischmann et al. [53] do not report on *Ocymyrmex Robustior* and *Cataglyphis Fortis* walking speeds. We discuss later these results in light of other species walking speeds during LWs and foraging walks.

#### Pirouette frequency decreases across successive walks

In their experiment, Müller and Wehner [55] counted pirouettes of *Ocymyrmex Robustior* within two meters of the nest during seven consecutive walks. In our simulations using the Müller layout, the number of pirouettes decreases sharply across successive walks (Fig. 5A), dropping from approximately 20 during the first walk to fewer than three from the third walk onward. A one-way ANOVA showed a strong effect of walk number on pirouette count (*F*_6,161_ = 12.53, *p <* 10*^−^*^10^), indicating that pirouette frequency differs significantly across walks, closely matching the experimental data from Müller and Wehner [55] (Fig. 5A).

**Fig 5.**
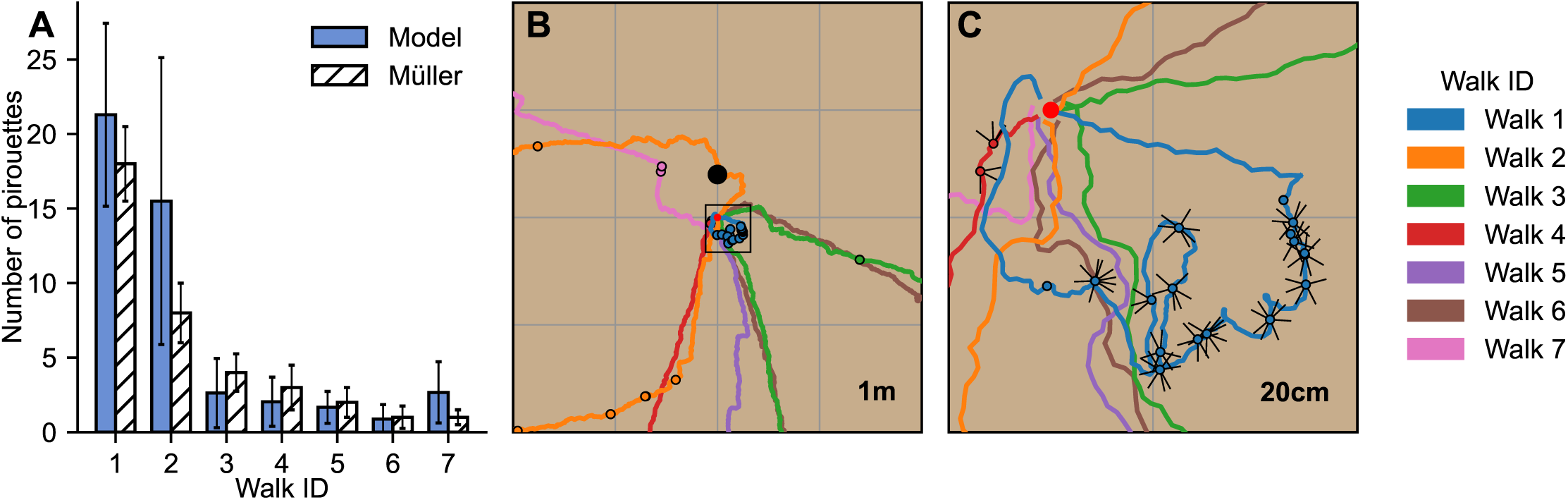
Pirouette frequency decreases across successive walks. **A:** The number of pirouettes decreases with walk ID in simulations of the Müller layout (blue bars), closely matching experimental data from *Ocymyrmex robustior* (white dashed bars; Müller and Wehner). Bars show mean ± 95% confidence intervals. **B:** Example trajectories from a single agent across seven successive walks, illustrating the progressive reduction in pirouette frequency and increasing trajectory regularity. First walk (blue) forms a very small loop confined within 30 cm of the nest with frequent pirouettes, the second and third walks (orange and green) extend to nearly 3 m from the nest with fewer pirouettes, and later walks (red, purple, brown, and pink) exceed 3 m and are straight foraging trajectories without pirouettes. The grey grid has one meter spacing; the nest is at (0,0), and the cylinder is shown as a black dot. Pirouette locations are shown as circles on the trajectories. Black square indicates close-up region shown in C. **C:** Close-up of the same walks as in B, showing snapshot acquisition during pirouettes (dots) and individual snapshots captured during the pirouettes (black arrows). The grey grid has a 20 cm spacing; Nest as a red disk.

Fig. 5B shows seven successive walks of a single agent while a zoomed-in view near the nest highlights the high density of pirouettes during the first LW (Fig. 5C). Notice the sharp decrease in frequency of pirouettes between the first and the second LW. This decrease is driven by an increase in visual knowledge as multiple snapshots are acquired in different directions during the pirouettes (black arrows, Fig. 5C). This in turn decreases the PI/VC error, lowering the probability of triggering a pirouette and thereby reducing pirouette count over a full LW.

#### Walks straighten with experience

Müller and Wehner [55] showed that path straightness increases across successive LWs (Fig. 6A). Simulations closely match this pattern, with a rapid increase in straightness during the early LWs. Four consecutive simulated walks from a single agent illustrate this progression (Fig. 6B). Early LWs form spirals around the nest, whereas later walks become increasingly directed. A one-way ANOVA on our simulated data showed a strong effect of walk number on straightness (*F*_6, 28_ = 47.02, *p <* 10*^−^*^30^). Path straightness rapidly increases during the first two LWs, followed by a plateau in subsequent walks. Post-hoc comparisons (Tukey HSD) showed that early walks (walks 1–2) were significantly less straight than later walks (walks 3–7; all *p <* 0.01), whereas differences among later walks were not significant.

**Fig 6.**
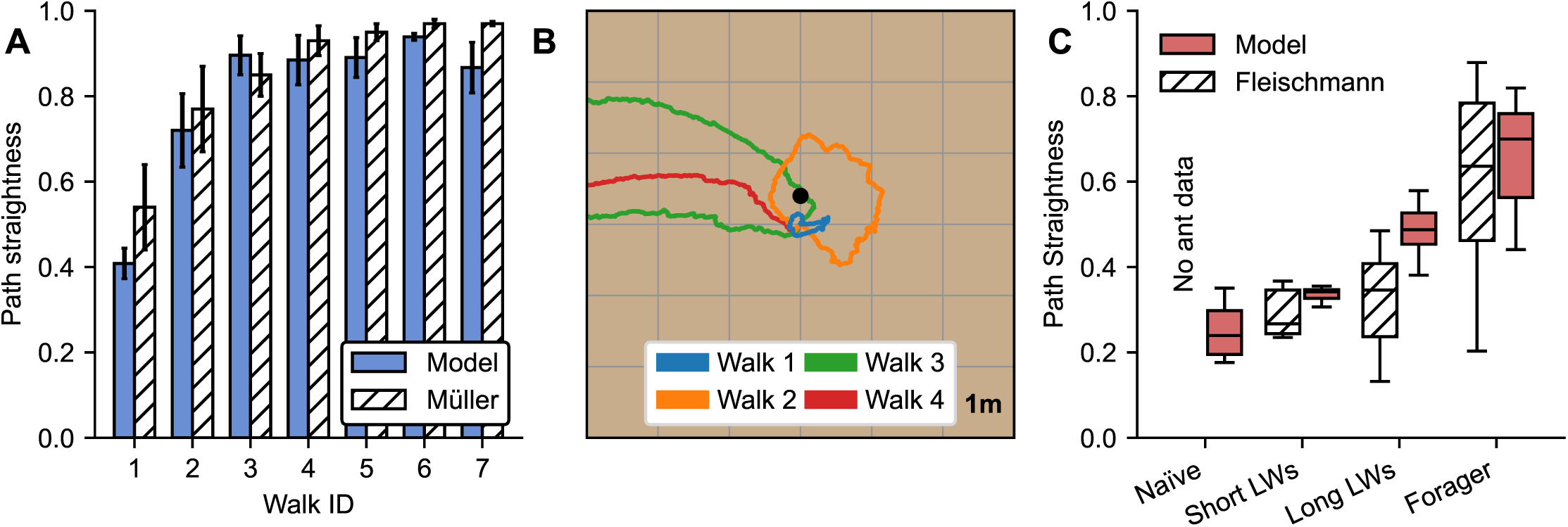
Walks straighten with experience. **A:** Path straightness increases across successive walks in simulations of the Müller layout (blue bars), closely matching experimental data (white dashed bars; Müller and Wehner [55]). Bars show mean ± 95% confidence intervals. **B:** Example trajectories from a single simulated agent across four successive walks in the Müller layout, illustrating progressive straightening. The first two walks (blue, orange) are LWs, while the third and fourth (green and red) are foraging walks. The grey grid has 1 m spacing; The small black dot represent the position of the cylinder at 40 cm north from the nest. **C:** Path straightness increases with experience in simulations of the Fleischmann layout (red) and matches experimental data (white dashed boxes; Fleischmann et al. [53]). Categories reflect increasing experience based on prior walk history: näıve, short LWs (*<* 0.7 m), long LWs (*<* 3 m), and foraging (*>* 3 m). Boxes show median and interquartile range; Whiskers extend to 1.5× the interquartile range; outliers are not shown.

A similar trend was reported by Fleischmann in *Cataglyphis fortis* (Fig. 6C). Instead of analyzing successive walks, they categorized ants into discrete experience groups based on prior exposure to the environment. These four categories included naïve individuals (no prior walks), ants that had performed short LWs (*<* 0.7 m), long LWs (*<* 3 m), and foraging ants (*>* 3 m). Our simulations closely reproduce Fleischmann’s data, with more experienced individuals producing straighter trajectories.

#### LWs expand with successive walks and accumulating experience

Fleischmann et al., [53] show example walks from desert ants in which walks become longer, cover larger areas, and extend further from the nest with increasing walk index (Fig. 7A). Our simulations reproduce this pattern, with successive LWs expanding markedly from one walk to the next (Fig. 7B,C). During the first four successive walks of our simulated agents we see expansion in distance from the nest (Fig. 7D), path length (Fig. 7E) and area (Fig. 7F), as confirmed by linear regression. Walks expand with walk ID across: Maximum distance from the nest (slope = 1.85, *R*^2^ = 0.28, *p* = 0.036), total path length (slope = 5.09, *R*^2^ = 0.42, *p* = 0.0068), and area covered (slope = 3.54, *R*^2^ = 0.39, *p* = 0.0099). This expansion emerges from the dynamics of *E*_PI_*_/_*_VC_. Increasing visual familiarity slows error accumulation and delays return to the nest. In parallel, increased walking speed and straighter trajectories promote the spatial expansion of LWs.

**Fig 7.**
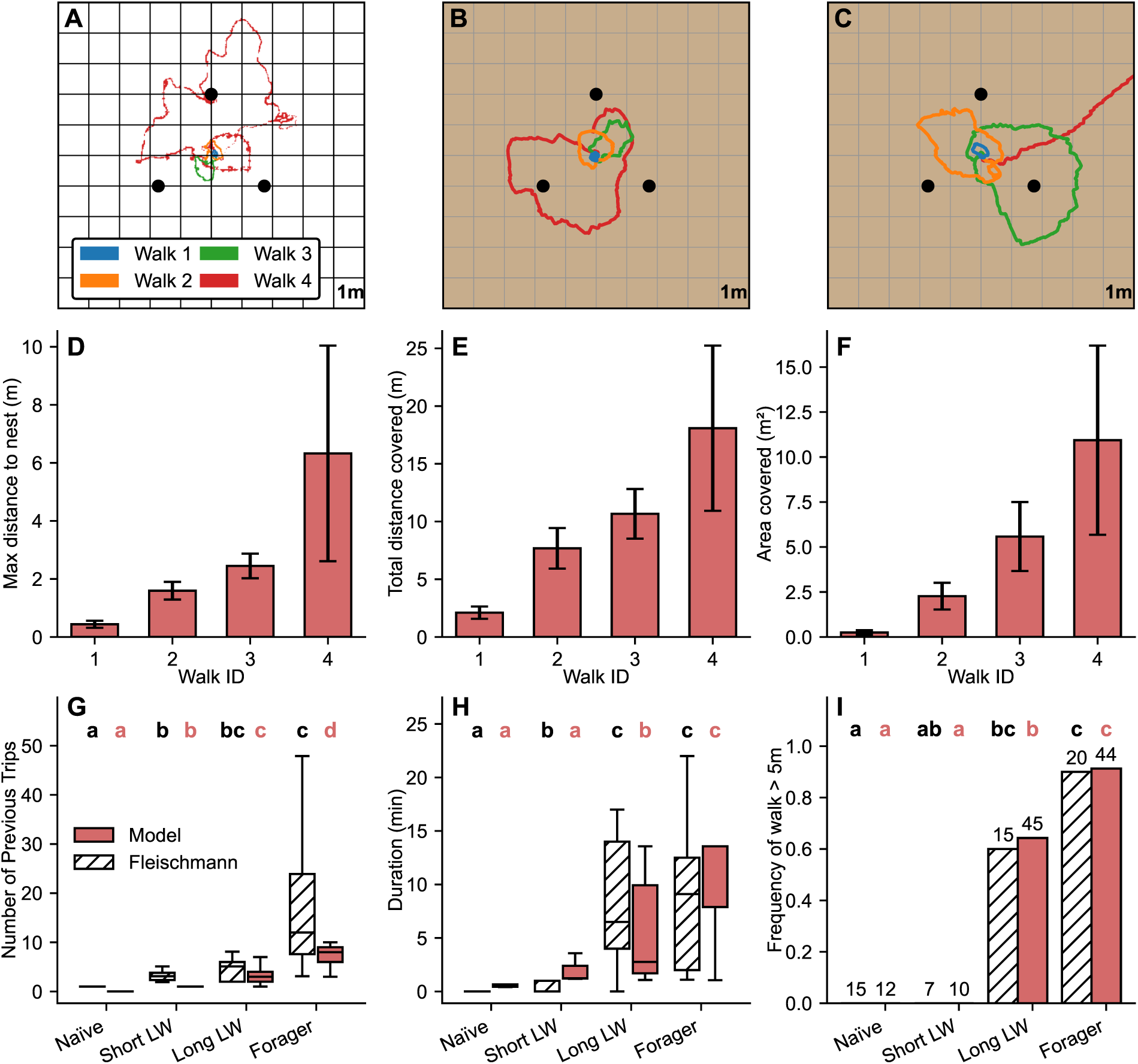
Walks expand with experience. **A:** Experimental trajectories from [53]. Blue: first LW of a näıve ant. Orange: second LW of an ant which had previously only performed LWs within 0.7 m of the nest). Green and red: third and fourth LWs of an ant which had previously performed LWs 0.7–3 m from the nest. The nest is at the center of the landmark array, surrounded by three black cylinders. Grid spacing is 1 m. **B:** First four successive walks of one of our simulated agent clearly showing the spatial expansion of LWs. The first walk (blue) is very short and remains within 30 cm of the nest. The second and third walks (orange and green) extend to ∼1.5 m. The fourth (red) is much larger but remains a LW (*<* 3 m). **C:** Another example of successive walks expanding for another simulated agent. The first three walks (blue, orange, green) are LWs, while the fourth (red) is a foraging trajectory (*>* 3 m; return phase not simulated). **D–F:** Walks expand across successive walks in simulations, as shown by multiple metrics. (**D**) Maximum distance from the nest, (**E**) total path length, and (**F**) area covered all increase across the first four walks. Error bars indicate 95% confidence intervals. **G:** Number of prior trips increases with experience categories (experimental data from [53] as white dashed boxes, model in red,). Boxes show median and interquartile range; whiskers extend to 1.5× the interquartile range; outliers are not shown. **H:** Walk duration increases across experience categories, with a marked increase between näıve ants and those that had made short LWs; and between those that had made longer LWs and foragers in both ants (dashed white) and the model (red). In addition to the progressive expansion of LWs, a foraging walk is triggered once agents exceed 3 m from the nest, adding 10 min to the walk (1 step = 200 ms). **I:** Proportion of walks extending beyond 5 m increases with experience; only experienced and foraging ants perform such long walks (dashed white bars). The model reproduces this pattern (red bars). Letters indicate statistical groupings based on pairwise comparisons (Kruskal–Wallis test or Fisher’s exact test, see text); groups sharing a letter are not significantly different. Numbers above bars indicate sample size.

As Fleischmann et al. [53] grouped ants into four categories of increasing experience rather than by successive walks, we next reproduce the authors’ original analyses and directly compare our model data with their experimental observations. In Fig. 7G, we see that in our model, higher experience categories are associated with an increasing number of previous trips (means: 0.0, 1.0, 3.43, 7.35; Kruskal–Wallis test: *p* = 1.23 × 10*^−^*^6^). Thus, as in Fleischmann et al. [53], we confirm that these four categories reflect a progression of experience. Pairwise comparisons show that all categories differ significantly from one another (all *p <* 0.05). Next, trip duration was analysed to quantify temporal expansion of LWs. In the simulations, walk duration increases with experience category (means: 0.62, 2.00, 5.27, 10.43 min; Kruskal–Wallis test: *p* = 1.97 × 10*^−^*^4^; Fig. 7H). The trend is consistent with Fleischmann et al. [53]. Pairwise comparisons show that näıve agents and those that had only performed short LWs do not differ (*p* = 0.114), whereas walk durations differ significantly between näıve agents and both those that had performed longer LWs (*p* = 0.0065) or transitioned to foraging behaviour (*p* = 0.0064). Similarly the walk durations of agents that had only performed longer LWs and foragers also differed significantly (*p* = 0.017). In our simulations, the timestep duration was set to 200 ms to match empirical data (Fig. 7H), and 1500 timesteps were simulated for each search.

Finally, the probability of leaving the vicinity of the nest was analysed as an indicator of expansion of LWs and the transition from LWs to foraging behaviour. We use a five meter threshold to match the analysis of Fleischmann et al [53]. In the simulations, the proportion of walks extending beyond five meters is 0% in näıve agents and in those that had only performed short LWs, increases to approximately 64% in agents that had previously made long LWs, and reaches 91% in experienced foragers (Fig. 7I). The proportion of ants leaving the five meters field differs significantly (Fisher’s exact test) between ants who had performed longer LWs and näıve ants (*p* = 0.0116) and from those who had only performed shorter LWs (*p* = 0.0179). Similarly, this proportion differs significantly between foragers and all other categories (näıve: *p* = 2.85 × 10*^−^*^6^; short LWs: *p* = 5.16 × 10*^−^*^5^; long LWs: *p* = 0.0179). Overall, the model reproduces the transition from short LWs to extended walks observed in Fleischmann et al., [53] experimental data.

Together, these analyses indicate a consistent spatio-temporal expansion of LWs over successive walks and accumulating experience, matching the qualitative trends reported for real ants.

#### Homing performance improves with experience

Fleischmann et al. [53] displaced zero-vector ants to a test field containing the same landmark array that surrounded the nest during previous LWs (Fig. 3B). Individually marked ants from each experience category were captured just before entering the nest, displaced to release points located three meters from the fictive nest position, and their search behaviour was recorded for five minutes. The authors analysed search accuracy (defined as the distance between the centre of the search density and the fictive nest) and homing success rate (whether the ant crossed the fictive nest position during the test). They reported a strong effect of experience on both measures. Näıve ants and those which had only performed short LWs showed inaccurate searches (Fig. 8I) centred on the release point rather than the nest [53] (Fig. 8D). In contrast, ants which had already performed long LWs or foraging walks exhibited more accurate searches (Fig. 8I), closely focused on the fictive nest position (Fig. 8H). Correspondingly, the proportion of ants reaching the fictive nest increased with experience. No näıve individuals or those who had only performed short LWs reached the fictive nest, whereas success rates increased to 33% after long LWs and 73% for experienced foragers (Fig. 8J).

**Fig 8.**
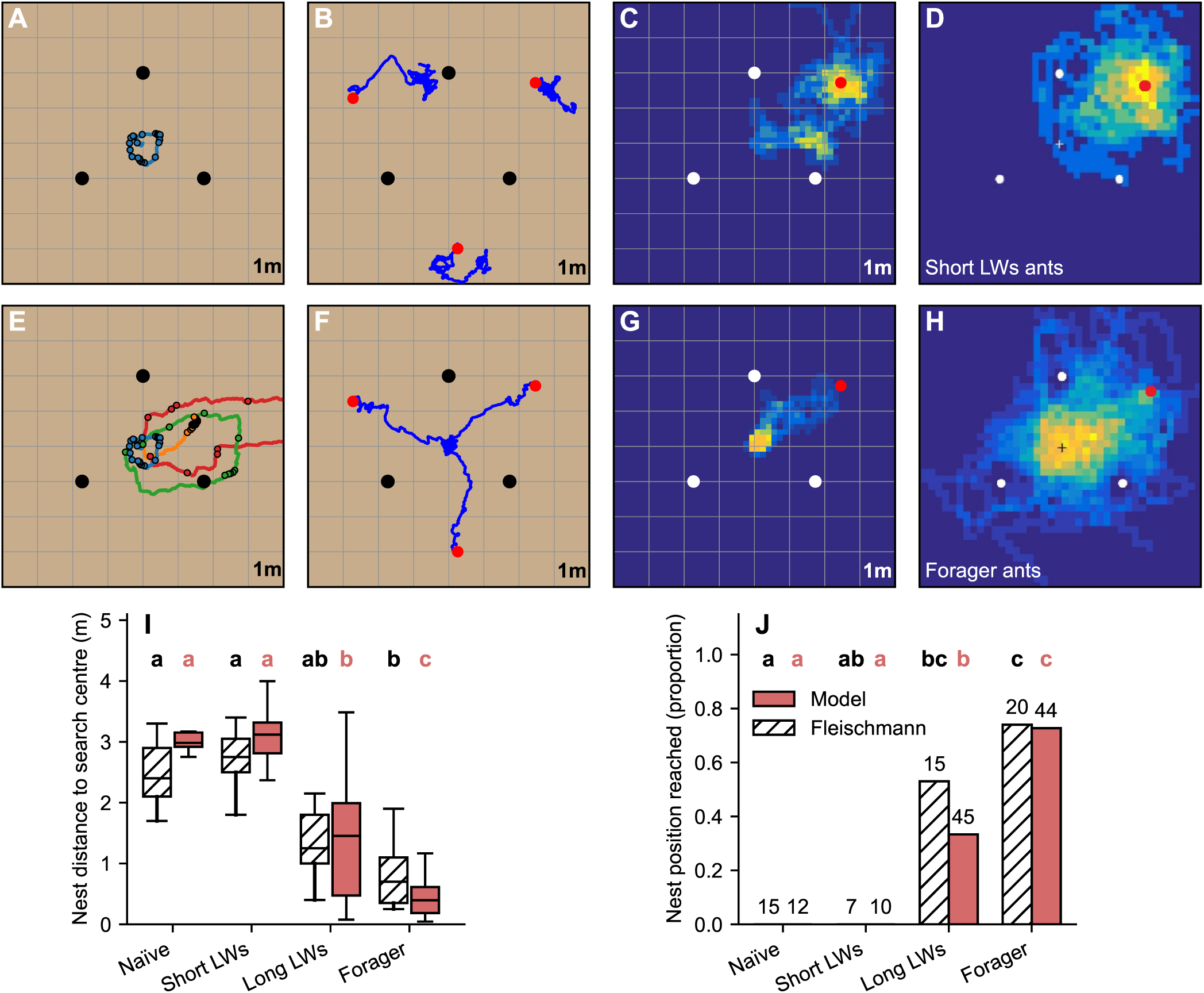
Homing performance improves with experience. **A** First and only LW of a category 2 agent. This short LW extends less than 0.7 m from the nest. The trajectory is shown in blue; pirouette locations are indicated by black circles. The three cylinders are shown as large black dots. Grid spacing is 1 m; the nest is at the center. **B** Three example trajectories of category 2 agents during visual-homing tests with zeroed PI vector. Agents fail to orient toward the nest. Release points are shown as red dots. **C** Density map of search locations for category 2 agents released at the same point. Searches are centred on the release point rather than the nest. **D** Experimental data from [53]: searches density map for category 2 ants (LWs *<* 0.7 m). **E** All LWs of a category 4 agent (blue, orange, green, red). **F** Four example trajectories of category 4 agents during visual-homing tests. Agents successfully reach the nest. **G** Density map of searches for category 4 agents. At this level of experience searches are centred on the nest. **H** Experimental data from [53]: search density map for forager ants. **I** Search accuracy for ants (dashed white boxes) and the model (red boxes) across experience levels. Accuracy is defined as the distance between the search center and the goal. Boxes show median and interquartile range; whiskers indicate the data range. **J** Proportion of ants crossing the fictive nest location when released 3 m away during tests [53], experimental data as dashed white bars, model results in red. A crossing is defined as approaching within 0.1 m of the nest. Letters above bars and boxplots indicate statistical groupings based on pairwise comparisons with Bonferroni–Holm correction; groups sharing a letter are not significantly different. Numbers above bars indicate the total number of walks (sample size) in each category.

Our model reproduces these trends quantitatively. Search accuracy differed significantly across experience levels (Kruskal–Wallis: *H*(3) = 70.43, *p <* 0.001; *n* = 12, 21, 45, 44 for increasing experience categories). Pairwise comparisons revealed that näıve agents and those which had only performed short LWs did not differ from each other (*p*_adj_ = 0.579), but both groups differed significantly from agents that had performed long LWs and from foragers. In addition, search accuracy improved significantly between agents which had performed long LWs and those which had begun foraging(Fig. 8I).

Homing success showed a similarly strong dependence on experience (pairwise Fisher’s exact tests with Holm correction). No agents in the näıve and short LWs groups reached the nest (0/12 and 0/21; Fig. 8J), whereas success rates increased to 33% (15/45) after long LWs and 73% (32/44) for foragers. Näıve and short LWs agents did not differ from each other (*p*_adj_ = 1.000), but näıve agents differed significantly from those that had preformed long LWs (*p*_adj_ = 0.052) and foragers (*p*_adj_ = 0.0001). Similarly agents which had only performed short LWs differed significantly from those which had performed long LWs (*p*_adj_ = 0.0045) and foragers (*p*_adj_ *<* 0.0001). Long LWs agents also differed significantly from foragers (*p*_adj_ = 0.0011).

Together, these results show that homing performance improves with accumulated experience. Näıve agents and those performing only short LWs do not reach the nest when released from a three meters distance (Fig. 8C), which is expected given that their experience is limited to the immediate vicinity of the nest (Fig. 8A). This does not imply that these agents are unable to home from shorter release distances. In contrast, both search accuracy and success rate are markedly higher in agents that have performed longer LWs (0.7–3 m from the nest) and improve further in experienced foragers (Fig. 8I-J).

#### Transition to foraging

The number of LWs performed before the first foraging trip appears to be broadly conserved across ant species and environmental contexts. Fleischmann et al. [53] reported that, in a largely featureless salt-pan environment with three artificial landmarks, *Cataglyphis fortis* performed between three and seven LWs before transitioning to foraging, whereas Jayatilaka et al. [58] found that, in a visually cluttered woodland environment, *Myrmecia croslandi* performed two to seven LWs (Fig. 9A), and Deeti et al. [63] reported that, in a semi-arid desert habitat, *Melophorus bagoti* performed three to seven LWs. In our reconstruction of Fleischmann et al.’s environment, simulated agents transitioned to foraging after two to seven LWs, matching the three to seven LWs reported for *Cataglyphis fortis* [53]. Using the same parameter set in Müller and Wehner’s environment, agents performed their first foraging walk (*>* 3, m from the nest) after only one to three LWs (Fig. 9A), a markedly lower range than the two to seven LWs reported across the literature [53, 58, 63]. M”uller and Wehner did not report the number of LWs preceding foraging. In our model, the transition to foraging occur at different rates depending on the structure and complexity of the environment.

**Fig 9.**
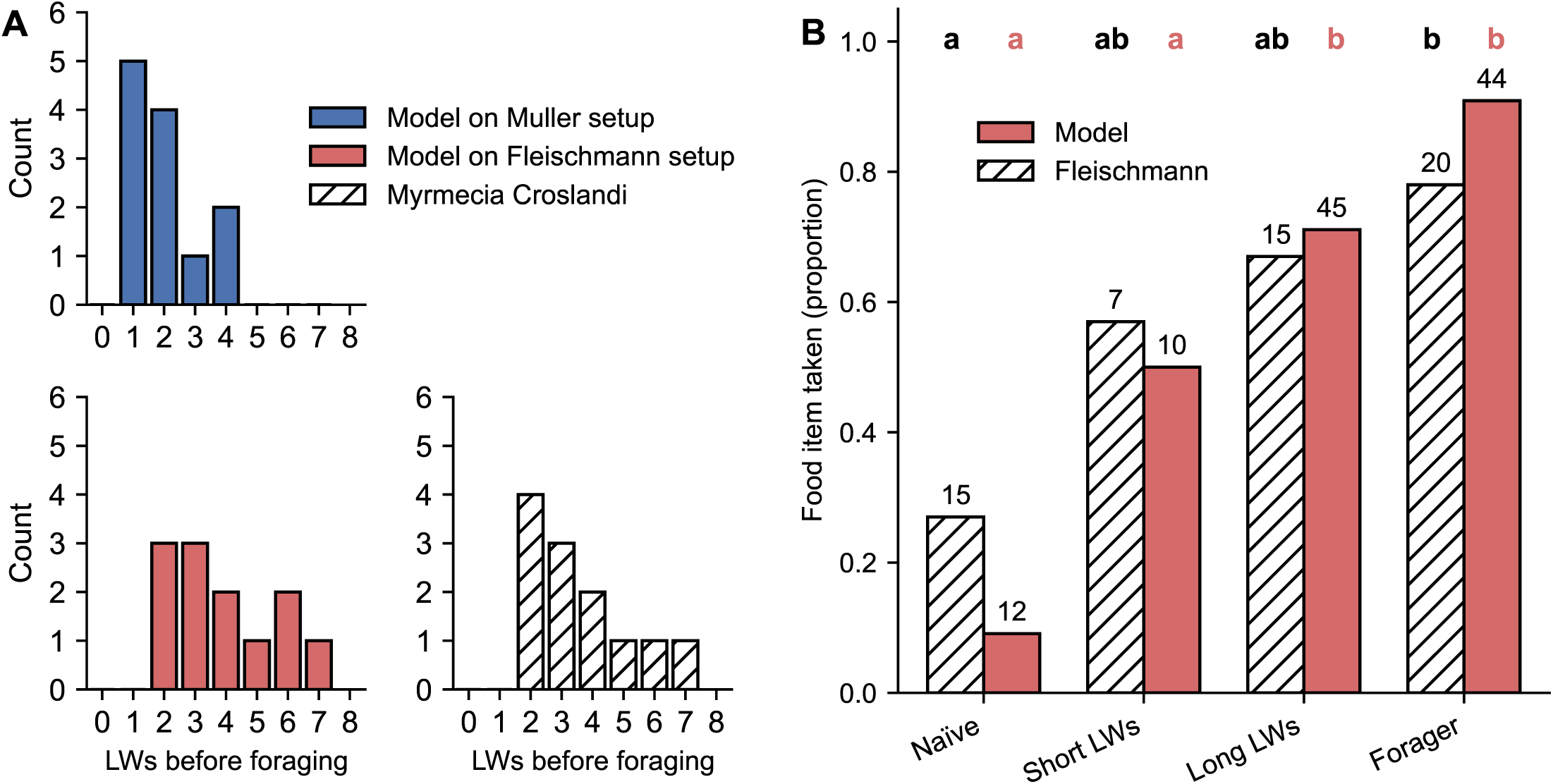
Transition to foraging. **(A)** Distribution of the number of LWs performed before the first foraging walk. In the model, agents in the Müller layout (blue) typically perform one to four LWs, whereas agents in the Fleischmann layout (red) perform two to seven LWs before the first foraging walk. Experimental data from *Myrmecia croslandi* (white dashed) show a similar distribution of two to seven LWs. **(B)** Proportion of ants of each category that picked up a bread crumb when presented and the derived behaviour from the model. Ants data as dashed white bars, model data as red bars. Letters above bars indicate statistical groupings based on pairwise Fisher’s exact test with Holm–Bonferroni correction; groups sharing a letter are not significantly different. Numbers above bars indicate the total number of walks (sample size) in each category.

Finally, Fleischmann et al. [53] found that the likelihood of ants picking up a food item increased with experience. Ants returning to the nest (after LWs or foraging walks) were captured and released at a location three meters from the nest position. When a cookie crumb was presented at the release point, näıve and short–LW ants rarely picked it up. In contrast, ants that had performed long LWs or were experienced foragers accepted the food in the majority of trials (Fig. 9B). We follow the same protocol in our simulations. In the model, food is picked at the release point if the recently accumulated *E*_PI_*_/_*_VC_ is above a fixed threshold. This mechanism was designed such that higher reliability of the visual compass promotes food acceptance. The model reproduces this pattern: food acceptance is low in näıve agents and those that had only performed short LWs, and high in agents that had performed long LWs or foraging trips (Fig. 9B). Statistical analysis using pairwise Fisher’s exact tests with Holm–Bonferroni correction confirms this pattern, with no statistically significant difference between näıve agents and those that had performed short LWs (*p*_adj_ = 0.127), and similarly no difference between agents that had performed long LWs and experienced foragers (*p*_adj_ = 0.087). However, there was a significant increase in food acceptance between näıve agents and both those that had performed long LWs (*p*_adj_ = 0.0013) and foragers (*p*_adj_ *<* 0.0001) as well as between agents which had only performed short LWs and foragers (*p*_adj_ = 0.028).

### Reoccurrence of learning after environmental change

Vermehren et al. [65] recorded experienced *Cataglyphis fortis* foragers leaving the nest after repeated training to a feeder, either with the nest surroundings unchanged (Control) or after introducing a novel black cylinder near the nest (Cylinder condition; Fig. 10A). The comparison tested whether a local visual change near the nest re-induces LW behaviours in experienced ants. We replicate this experiment using two arenas: a 1 m diameter arena without a cylinder (Control; Fig. 10B) and an identical arena with an added black cylinder (7.5 cm diameter, 11 cm high) placed 20 cm from the nest (Cylinder; Fig. 10A,C). Each agent was first trained for six walks in the no-cylinder configuration and, by this point, all agents had begun foraging (going *>* 3 m from the nest). We then separated the experienced agents into two groups, and recorded their behaviour during a full outward walk either without (Control) or with the added cylinder (Cylinder).

**Fig 10.**
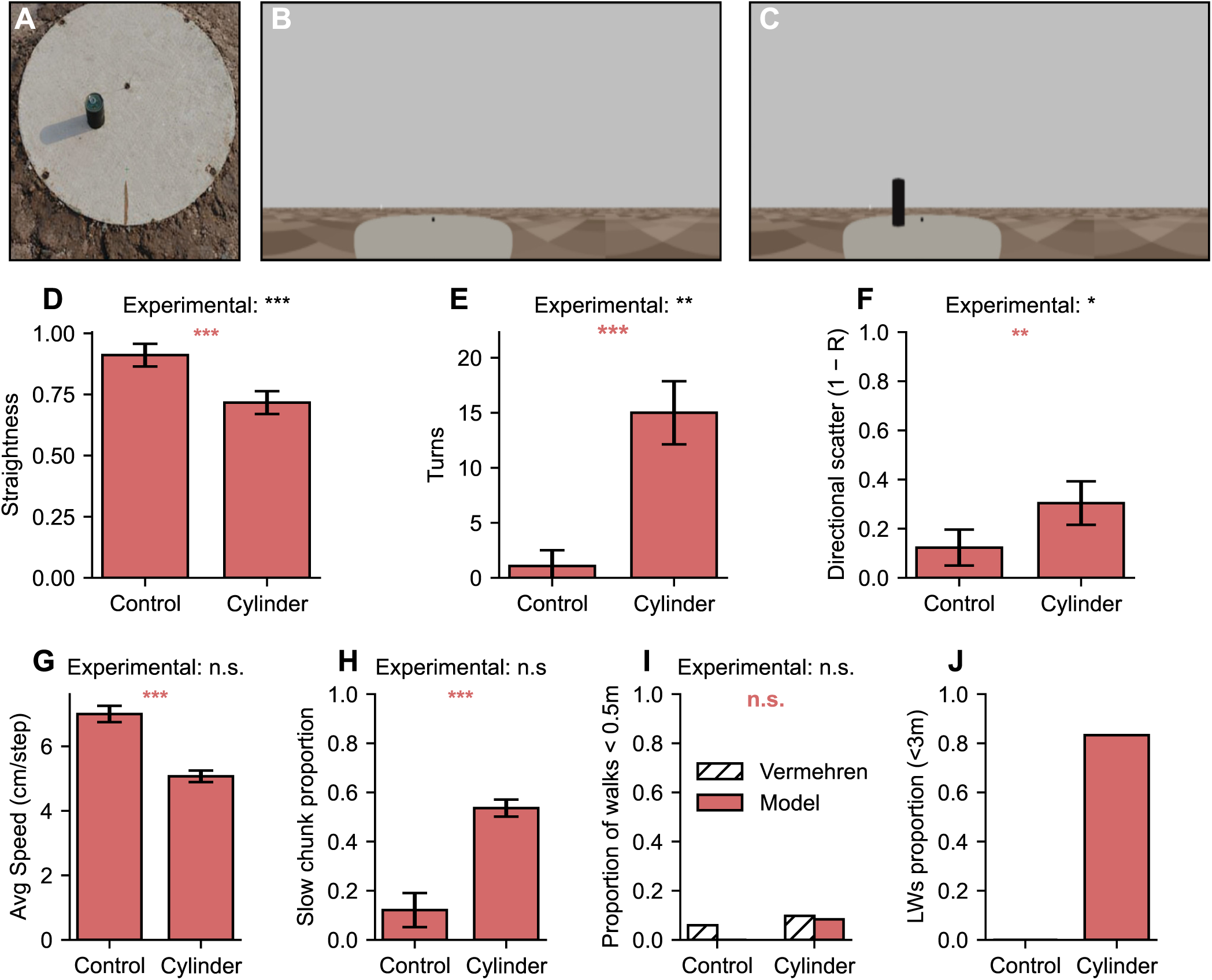
Reoccurrence of learning after environmental change. **A:** Original experimental layout (Vermerhen et al.). **B:** Agent view of the reconstructed control environment used for training, with corresponding 360° panoramic view taken 1 m from the nest. **C:** Agent view of the reconstructed environment with the cylinder introduced, after six prior walks using the control setup. **D:** Path straightness during outward walks. Straightness is significantly lower in the cylinder condition than in control. Error bars indicate 95% confidence intervals. **E:** Pirouette count per walk. Adding the cylinder triggers pirouettes, with a strong increase in turn frequency. **F:** Heading-direction scatter. Directional scatter is significantly higher in the cylinder condition, while mean heading direction remains broadly comparable. **G:** Mean walking speed during outward walks. In simulations, speed is significantly lower in the cylinder condition, but not in experimental data. **H:** Proportion of slow walking segments. In simulations, the proportion is higher in the cylinder condition, although experimental data report no significant difference. **I:** Proportion of “aborted walks” staying within 0.5 m of the nest. Most walks extend farther than 0.5 m from the nest. Abort rates do not differ significantly between conditions in either the model or experimental data. **J:** Proportion of LWs relative to foraging walks (simulated data only). There is a large proportion of LWs in the Cylinder group and no LWs in the Control group.

Vermehren et al. reported a significant reduction in straightness when a novel cylinder was placed near the nest. Our model reproduces this effect: path straightness was significantly lower in the Cylinder condition than in Control (Mann–Whitney *U* = 137.0, *p* = 1.96 × 10*^−^*^4^; see Fig. 10D). Additionally, Vermehren et al. showed that nest-aligned segments and the occurrence of nest-directed turns were higher in the Cylinder condition than in the Control condition. In our data, we quantify scans using a single measure: the number of pirouettes per walk. Pirouette counts were strongly elevated in the Cylinder condition relative to Control (Mann–Whitney U = 0.50, *p* = 3.31 × 10*^−^*^5^; ∼15 pirouettes vs ∼1 pirouettes in control; *n* = 12 per group, Fig. 10E), matching the results reported by Vermehren et al.

Following Vermehren et al., we tested whether the cylinder made outward walks less directed, as expected during exploration. Mean heading was unchanged between Control and Cylinder conditions (Control: 177.04*^◦^*; Cylinder: 180.57*^◦^*; Watson–Williams test: *F* = 0.159, *p* = 0.6905). However, heading scatter, quantified as 1 − *R* where *R* is the mean resultant length, increased with the cylinder (1 − *R* = 0.304 ± 0.088, *N* = 129) compared with Control (1 − *R* = 0.123 ± 0.073, *N* = 77), and this difference was significant (permutation test on *R*: Δ*R* = 0.181, *p <* 0.0001). Thus, like Vermehren et al., we found no change in average orientation but increased heading scatter, consistent with more exploratory walks.

Vermehren et al. also analysed walking speed by dividing outward paths into fixed two centimeter segments. They reported no significant difference between Control and Cylinder conditions. In contrast, our model shows a significant reduction in average speed in the Cylinder condition with ∼ 5.2 vs ∼ 6.7 cm/step (Mann–Whitney *U* = 144.0, *p* = 3.66 × 10*^−^*^5^; Fig. 10G). Vermehren et al. defined a ‘slow-walking’ segment as one which ants traversed at less than 0.06 m s*^−^*^1^) and, using this criterion, they found no difference in the overall proportion of slow segments between conditions, whereas our model exhibits a marked increase in the proportion of slow-walking chunks in the Cylinder condition with ∼ 0.51 vs ∼ 0.15 (*χ*^2^ = 158.63, *p* = 2.25 × 10*^−^*^36^; Fig. 10H). We found, however, that the outcome of this analysis depended on the chosen speed threshold.

Finally, Vermehren et al. analysed whether a novel landmark near the nest disrupted the ants’ departure from the nest. They quantified this by measuring how many ants failed to leave the platform (0.5 m radius) after emerging from the nest, treating such early returns as aborted foraging attempts. Using this same definition, they reported no significant difference in abort rates between Control and Cylinder groups. We replicated this analysis and similarly found no significant effect at the 0.5 m threshold, with 1/12 walks aborted in the Cylinder group and 0/12 in the Control group Fig. 10I).

Overall, the results in this section demonstrate that our model reproduces the behaviours seen in ants, where introducing a novel landmark near the nest induces ‘re-learning’ behaviours and even LWs.

## Discussion

### The active learning model

#### The active learning model captures the observed dynamics of learning walks

We showed that a single error signal can control LW geometry and dynamics, reproducing key behaviours of desert ants: expansion and straightening of LWs, a declining number of pirouettes, the transition to foraging, and the reappearance of LWs after panorama changes. Furthermore, the model remains robust under biologically realistic levels of compass noise.

The model reproduces the characteristic LW behaviours reported by Müller and Wehner [55] including early spiralling trajectories progressively straightening into foraging walks. Spiralling likely serves a dual function during LWs. First, it promotes sampling across many directions around the nest. Second, it limits drift away from the nest and prevents premature transition to foraging while the VC is still unreliable. Similarly, the model reproduces the progressive disappearance of pirouettes with learning reported by Müller and Wehner [55]. In our model, both the spiralling behaviour and the probability of triggering a pirouettes are function of the *E*_PI_*_/_*_VC_ signal and the disappearance of both pirouettes and spiralling is simply explained by learning of the VC, which leads to decreasing *E*_PI_*_/_*_VC_ (Fig. 4). Similarly, as *E*_PI_*_/_*_VC_ decreases with experience, walking speed increases and walks progressively straighten, leading to the characteristic LW expansion described by Fleischmann et al. [53]. Our model also reproduces the food-picking behaviour reported by Fleischmann et al. [53] where experienced foragers reliably pick up bread but näıve ants rarely do. While this pattern *could* arise from a simple history-dependent rule, it can be reproduced with our model by simply gating bread-picking with the same *E*_PI_*_/_*_VC_ signal used to regulates LW geometry. Through this lens, bread picking therefore just constitutes another behavioural marker of the transition from learning to exploitation. A prediction of this is that altering the visual panorama should suppress food acceptance in experienced ants.

Our model also reproduces the gradual improvement in visual homing performances with experience [53]: as LW and experience accumulate, agents are increasingly likely to cross the nest position during visual homing trials. This result demonstrate that visual landmark information acquired during successive LWs is integrated gradually, leading to progressively more accurate nest localisation and a larger catchment area. This is useful as LWs become longer and PI error increase, especially after long foraging walks. For example, after a 50 meter walk, Huber et al. [19] reported that the mean PI error relative to the true nest position is ∼ 7.5 m and increases to ∼ 14 m after a 150m walk. In such cases, the error magnitude of PI may still exceed the visual catchment area so visual learning may not entirely eliminate the need for systematic searches. However, visual learning could still provide a broad visual attractor that accelerates systematic search and provide a more reliable signal than olfactory plumes emanating from the nest, which may be sensitive to wind or in some cases (e.g. CO_2_) are not specific to a colony, leading ants to mistakenly enter the nest of a foreign colony [47].

Finally, our model replicates the key findings of Vermehren et al. [65], showing that changes in the visual panorama trigger re-learning behaviour, exploratory trajectories, and, in some cases, full LWs in experienced foragers. In their experiments, a novel landmark placed along the habitual route increased heading dispersion, reduced path straightness, and triggered nest-facing saccades, while walking speed remained unaffected. In our model, a panorama change produces qualitatively similar effects: trajectories become less straight, heading disperse, pirouettes reappear, and the proportion of LWs is higher than in the control condition. Unlike Vermehren et al. [65], we observe a clear reduction in walking speed following panorama change. Reduced walking speed during LW is a well-established signature in both ants [58] and bees [2], making the absence of a speed reduction reported by Vermehren et al. [65] somewhat surprising. Furthermore, the increase in walking speed we observed in our model is in line with that reported by Jayatilaka et al. [58], who record mean walking speeds of ∼ 12.7 mm/s during LWs and ∼ 25.2 mm/s during foraging.

#### Knowing when to learn

LW are costly as, during learning, ants do not forage and remain exposed to risks such as desiccation and predation away from the nest. Learning must therefore be brief, yet sufficient to correct subsequent PI drift and ensure reliable homing. By dynamically triggering LWs based on the instantaneous error (*E*_PI_*_/_*_VC_), our model parsimoniously balances these competing concerns, only triggering learning behaviours when visual guidance is unreliable. As the agreement between PI and vision improves, learning behaviours and LWs naturally vanish, but immediately resume if the panorama changes beyond recognition. Most importantly, this makes LWs adaptive to the specific environment faced by the agent – some environments might be easier to learn than others. For example, in our reconstruction of the environment used by Müller and Wehner [55] which only features a single black cylinder, agents transition to foraging much more quickly (one to four LWs) than they do in the environment with three cylinders used by Fleischmann et al. [53] (two to seven LWs). This suggests that learning speed may indeed by proportional to visual complexity and that our model can adaptively stop LWs after sufficient learning has been performed. This prevents both premature foraging and unnecessary or redundant learning.

In our model, learning reoccurs when the visual compass fails to predict the nest direction, but in some cases, for example in featureless or symmetric scenes, the panorama may not provide useful information for locating the nest. Interestingly, *Cataglyphis Velox* ants reduce the amplitude and regularity of their oscillations when visual scenes lack informative structure, suggesting the use of simple heuristics to determine whether a scene is worth exploring [75].

#### How could the model fit with known neuroanatomy

Anatomical evidence summarised by Heinze et al. [16], indicates that, in the fan-shaped body (FB) of the central complex, pathways carrying PI information from the protocerebral bridge via CPU4 columnar neurons converge with Mushroom Body Output Neuron (MBON) axons carrying learned visual information. This anatomical arrangement places PI-derived and visually derived familiarity signals within the same neuropile, making the computation of the error signals required by our model anatomically plausible. Our model therefore predicts that this signal should be identifiable in the FB; its discovery would provide clear physiological support for the model. Additionally, the eight MBONs required to implement the visual compass remains biologically plausible, given that Drosophila have 34 MBONs [76], honeybees around 400 [77], with ants likely to fall between these values.

#### Clawed walks (partial relearning walks)

Jayatilaka et al. [58] observed that experienced foragers sometimes perform short spiralling segments near the nest before starting a straighter foraging walk in the opposite direction. Jayatilaka et al. interpreted the as short learning walk segments to learn snapshots which prevent overshooting the nest during subsequent homing. A similar behaviour can be observed in our simulations. This behaviour emerges in our model because both PI and VC uncertainty is high near the nest, leading to high *E*_PI_*_/_*_VC_ which triggers spiralling. Our model suggests that these “clawed walks” could be a byproduct of using the mismatch between PI and visual cue to modulate exploration behaviour, rather than an adaptive, distinct behavioural mode to prevent overshooting the nest (see SI Fig. 11 for details).

#### Prediction: PI–vision conflict

Our model predicts that inducing a mismatch between PI and visual estimates of nest direction should reactivate learning behaviours. We would expect the magnitude of this response to scale with the angular offset between the two cues and we provide a testable protocol with predictions in the Supplementary Information section (SI Fig. 12 for details).

#### Generalisation to other solitary central-place foragers

The mismatch mechanism we presented here could apply to other ants species and flying hymenopterans such as bees and wasps, whose orientation flights share structural features with ant LWs. PI and visual compass are also available to these taxa, and may interact similarly to scaffold visual learning.

### The visual compass model

Finally, we discuss why we chose to use our own variant of the VC rather than the classical version [39] or a neural implementation of it [50].

#### Nest-oriented or multi-directional snapshots

Multiple fixations at different orientations have been observed in multiple ant species. For example, Fleischmann et al. [59] reported that *Cataglyphis noda* made 4 ± 3 fixations during each pirouette. Consistent with this, detailed frame-by-frame analyses of LWs in *Myrmecia croslandi* and *Melophorus bagoti* revealed that neither overall gaze directions nor low-angular-velocity “fixations” exhibit a peaked nest-centred (or anti-nest) distribution; instead, gaze directions broadly and relatively homogeneously cover the full azimuthal range around the nest [73]. Our implementation provides a functional hypothesis for this behaviour as, by learning multiple snapshots at different orientations, rIDFs can be calculated internally, in parallel across MBONs.

#### Sampling during LWs

In our active learning model, *E*_PI_*_/_*_VC_ must be computed continuously to regulate the balance between exploration and exploitation. In the classical VC framework, inference is tightly coupled to sampling behaviour meaning that, in order to predict the direction of the nest, agents must perform physical scans that overlap the nest direction. However, unlike during homing, ants are typically not oriented toward the nest during the majority of a LW so it is not clear to us how nest direction could be continuously estimated using the classical VC framework. Our model provides continuous estimates of the nest direction by calculating nest direction using multiple MBONs However, in order to learn these memories, scanning behaviour is required during learning. In this work we only model pirouettes, but other types of scanning behaviour have been observed during LWs such as voltes [59] or the large ∼ 180*^◦^* continuous oscillations observed in *Myrmecia croslandi*, *Melophorus bagoti*, and *Cataglyphis nodus* [54, 58]. Interestingly, oscillation amplitude is higher during LWs than during foraging walks [54, 58], suggesting a similar functional role as pirouette in visual learning. The pirouettes in our model could probably be replaced with oscillations, which amplitude could depend on *E*_PI_*_/_*_V_.

#### Sampling during foraging

The decoupling of inference and scanning become critical during foraging. Unlike during LWs, experienced foragers typically leave the nest hastily along relatively straight trajectories without significant oscillations [58]. Yet, they remain sensitive to changes in the visual landscape, which can re-induce learning behaviours such as pirouettes. In the classic VC framework, experienced foragers would lose their ability to compute *E*_PI_*_/_*_VC_ while they don’t face the nest, hence hindering their ability to compute the reliability of their VC, at least within the scope of our active learning model. In this context, our visual compass model provides a more parsimonious solution by allowing directional predictions to be computed independently of the agent’s heading. Our VC and PI models both generate precise egocentric predictions of nest direction, without the need for physical scan and nest fixations, which facilitates uninterrupted computation of *E*_PI_*_/_*_VC_. Similarly to the model of VC proposed by Wystrach et al [73], our VC model transfers the burden of performing physical rotational scans from inference-time to training-time. While physical rotation occurs during learning through pirouettes, during inference, the eight parallel MBONs compute an instantaneous scan in-silico.

#### Limitations

Our model cannot explain several phenomena observed in ants. First, high-speed analyses have shown that nest-oriented fixations are significantly longer than non–nest-oriented fixations [59]. Moreover, ants and flying insects are notorious for fixating the nest during pirouettes and learning flights [38, 55, 60, 61]. However, our model fails to explain the functional purpose of this bias towards nest-directed fixations. Our VC and active learning models function without any nest-oriented bias during pirouettes, as the full panorama is uniformly sampled (SI Fig. 13E-F). One possibility is that multiple VC strategies coexist in the insect brain, which could lead to the ants sampling the full panorama, while showing a bias toward nest-oriented fixations. Alternatively, during homing, ants mainly face toward the nest, and biasing learning toward that direction might be adaptive.

#### Conclusion

In short this visual compass model gives another perspective on insect navigation compared to the classic model [39], opponent models which capture nest and anti-nest views [78, 79] and lateralized models of the visual compass [51, 73, 80]. While it doesn’t explain the bias of insects to fixate the nest during LWs, it enables precise predictions without scanning, decoupling inference from gaze direction. It fits with our active learning model and gives a functional role to the non-nest oriented fixations observed in desert ants [59] during pirouettes.

## Materials and methods

### Antworld renderer

Panoramic views of the environment were generated using the *Antworld* rendering pipeline from the publicly available Brains-on-Board robotics library [43, 81], as in our previous work [82]. *Antworld* renders RGB panoramas of 3D scenes at selected positions and orientations. Here, the virtual camera was positioned five centimeters above the ground – slightly higher than a real ant’s viewpoint – to avoid artefacts where the camera goes through the ground plane. Views were rendered with a 360° horizontal and 75° vertical field of view.

### Experimental layouts and protocols

We reconstructed the visual environments of three published experiments on desert ant LWs [53, 55, 65]. For each study, we recreated a minimal 3D arena directly from the experimental descriptions. The arenas replicate a sandy, largely featureless desert environment with a small number of salient landmarks arranged according to the original configurations. Each scene was procedurally generated in Python and exported as a textured .obj mesh, then rendered in *Antworld* to generate panoramic visual input for the model.

### Müller & Wehner (2010) [55]

A 40 m × 40 m flat sand plane was generated to reproduce the uniform salt-pan surface described by Müller & Wehner. A 11 cm diameter, 0.3 m tall black cylinder was positioned 0.4 m north of the nest, matching the layout of the original experiment (Fig. 3A,B for visualization). The sand surface colour was set to RGB ≈ (181, 162, 119), corresponding to the average colour of a patch sampled from a photograph of the nest surroundings provided by Fleischmann et al. [59], approximating the natural tones of the Namibian desert.

Seven consecutive walks were simulated for each agents, starting from the nest entrance. The initial azimuth of the agent was randomly set at the beginning of each walk and each trial lasted up to 1000 steps or ended when the agent returned within 30 cm of the nest. From this point, the agent was guided along a straight path to the nest entrance, and the walk was terminated once it was within 1 cm of the nest. The walks were recorded without intervention, but the simulation was stopped if the agent went farther than 5 m from the nest. Pirouettes were recorded up to 2 m from the nest. Straightness was quantified as the ratio between the beeline distance from the nest to the farthest point reached and the total distance walked up to that point:

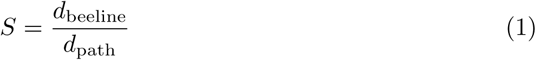

where *d*_beeline_ is the Euclidean distance between the nest and the maximum distance reached, and *d*_path_ is the cumulative path length traveled until that point.

### Fleischmann et al. (2016) [53]

A 40 m × 40 m flat sand plane was generated as above. Following Fleischmann et al. [53], three equidistant black cylinders (diameter 22 cm, height 38 cm) were placed 2 m from the nest entrance at azimuths 90°, 210°, and 330°. See Fig 3 for visualization. The nest was represented by a small central hole (diameter 3 cm). A 10 m × 10 m white grid (1 m spacing, 3 cm lines) was embedded in the sand, reproducing the calibration grid visible in the original recordings, which provided positional scale and reference, to track ant trajectories, but serving no functional role in our simulations. Fleischmann et al. assigned ants to four experience categories based on the maximum distance reached from the nest during their previous walks:

- **Category 1 (Näıve):** first outdoor appearance and previous excursions confined within *<* 0.3 m of the nest.
- **Category 2 (Short learning walks, SLW):** one or more LWs, never exceeding 0.7 m.
- **Category 3 (Long learning walks, LLW):** Previous LWs extending beyond 0.7 m but not exceeding three meters from the nest.
- **Category 4 (Foragers):** Previous foraging trips occurred, exceeding three meters from the nest.

Our protocol followed the same procedure as in the Müller and Wehner replication, with the following differences. First, for each agent, a variable number of LWs was simulated to reproduce the variability in the number of walks observed in the original study. The number of LWs per agent was fixed to 9. Each LW lasted a maximum of 2000 steps and ended if the agents returned to the nest, following the same protocol as the Müller & Wehner experiment.

In addition, agents were submitted to visual homing tests. During each test, agents were released at a random location 3 m from the nest and allowed to navigate for up to 1500 steps. Search patterns were recorded during the final 1000 steps of each test. Search accuracy was computed as the distance between the centroid of the search trajectory and the fictive nest position. Homing was considered successful if the agent approached within ten centimeters of the nest. In the simulations, the timestep duration was set to 200 ms to match empirical data (Fig. 7E). If the agent crossed a distance superior to five meters from the nest, fifteen minutes (4500 steps) were artificially added to the trajectory to simulate a foraging walk after escaping the nest vicinity. This closely matches the empirical data (Fig. 8E).

Unlike Fleischmann et al. [53], who tested each ant only once after performing a random number of walks and then removed from the experiment, tests in our simulations were interspersed with normal (LW or foraging) walks. Each agent was tested once without prior experience and again after every subsequent LW. To prevent early tests from influencing later behaviour, learning was artificially disabled during all tests.

### Vermehren et al. (2020) [65]

A 40 m × 40 m flat sand plane was generated as above and we recreated the four experimental contexts described by Vermehren et al. . The default layout was composed of a 1 m diameter beige platform and the 3 cm nest hole at the origin of the arena. These objects contrasted with the brown surrounding sand RGB ∼ (214, 199, 156), sampled from the average of a patch of a photo provided in the original paper. Three additional contexts were created, where a single 7.5 cm black cylinder was positioned 20 cm north, east, or west of the nest. See Fig 3 for visualization. The protocol followed the same procedure as in the Müller and Wehner replication. All agents first performed five LWs in the default layout. Agents were then assigned to one of two groups: in the landmark condition, a single cylinder was added to the scene in one of three possible configurations, whereas in the control condition the scene remained unchanged. In both cases, the behaviour of the agent was recorded during the subsequent walk.

### Path Integration

We implemented a Path Integration (PI) model, inspired by the model of the insect Central Complex (CX) developed by Stone et al. [74]. We used *N* = 8 directional columns, reflecting the eightfold columnar organization of the protocerebral bridge [16, 74]. The PI memory is encoded by eight CPU4 neurons, maintained in an allocentric reference frame, with heading represented relative to a fixed external compass rather than the agent’s body axis. The agent’s heading is encoded by eight TB1 neurons, which encode head-direction in the CX [16, 74], with cosine tuning:

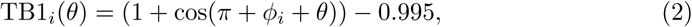

where *ϕ_i_* are the preferred directions of the columns. PI is noisy and accumulates error, which can make the signal used to orient snapshots and nest-directed fixations imprecise. Muller et al. [55] reported a mean angular deviation of 1.9*^◦^* (SD ∼23.5*^◦^*, *n* = 163) in *O. robustior*, and Fleischmann et al. [59] found comparable dispersion near the nest in *C. noda* (mean 7.5*^◦^*, 95% CI −5.3*^◦^*–20.2*^◦^*, *n* = 83) and *C. aenescens* (3.5*^◦^*, 95% CI −7.5*^◦^*–14.4*^◦^*, *n* = 11). To reproduce these error levels, in all experiments, we added noise to the compass signal, a standard approach to model noise in PI models [69]. At each step, the agent’s heading *θ* is corrupted by zero mean Gaussian noise (*σ_θ_* = 50*^◦^*).

Forward speed is encoded by a TN1 signal [16, 74] and TB1 and TN1 are combined to update the CPU4 memory:

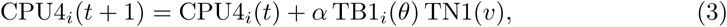

where *θ* is the noisy heading, *v* denotes forward speed, *α* = 0.05 is the memory gain, and no odometric noise is applied (*σ_v_* = 0). The CPU4 memory was initialized to zero and updated by linear accumulation, with no rectification, clipping, nonlinear activation.

The nest azimuth is decoded by population-vector averaging over the CPU4 memory:

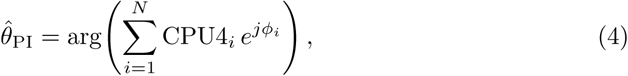

where *ϕ_i_*are the column preferred directions (offset by 45*^◦^* to align with the eight cardinal directions).

### Vision model

Panoramic RGB views rendered with the *Antworld* pipeline were processed through minimal preprocessing steps that are common in the insect navigation literature: isolation of the green pathways, lateral inhibition and downsampling [39, 69, 82]. The green channel **I***_g_* was extracted, downsampled to 36×36 px and filtered by a lateral inhibition kernel using a center-surround mechanism. These operations greatly increases the performance and capacity of the MB model [83] and have been widely reported in the literature as means of enhancing spatial and temporal contrast and supporting efficient encoding [84–86].

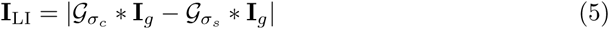

where G*_σ_*denotes a Gaussian kernel with standard deviation *σ*, and ∗ denotes a 2D convolution.

The resulting feature map was downsampled to 10×10 px and flattened into a 100-dimensional vector used as visual input to the memory model. While this simplified representation does not aim to capture the actual input to the ants’ Visual Projection Neurons (VPNs), its dimensionality is consistent with the order of magnitude of VPNs reported in insects (e.g. ∼400 in honeybees [87]), here we conservatively use a lower number of VPNs.

### The Visual Compass model

We implement the visual compass as an MB-inspired network with a ring of eight directionally tuned MBONs. This architecture builds on the lateralized model of Wystrach et al. [51, 73], in which visual memories are distributed between MBONs encoding whether the goal lies to the left or right of the agent’s current heading. Here, we extend this principle to eight MBONs, each associated with a preferred egocentric nest azimuth, thereby increasing the angular resolution of visual-compass predictions and matching the eight-column layout of the CX. This allows an error signal to be computed between the PI estimate and the visual-compass prediction. The vector of 100 VPNs carrying panoramic visual input information is projected onto a population of *n*_KC_ = 20,000 Kenyon cells (KCs) through fixed, sparse, random connectivity: each KC receives input from exactly 5 randomly selected input units [88]. KC activation is computed as the linear sum of its inputs, followed by sparsification so that only the top 1% of KCs are active. The resulting KC output is binary: selected KCs are assigned an activity of 1, while all remaining KCs are assigned an activity of 0. We implement a ring of 8 directionally tuned MBONs with preferred egocentric nest azimuths at {−157.5*^◦^,* −112.5*^◦^,* −67.5*^◦^,* −22.5*^◦^,* 22.5*^◦^,* 67.5*^◦^,* 112.5*^◦^,* 157.5*^◦^*}. Each MBON receives input from the full KC population through independent synaptic weight vectors initialised with random uniform weights between 0 and 1.

Plasticity occurs at the KC–MBON synapses. During learning, the MBON whose preferred direction is closest (in circular angular distance) to the current egocentric nest azimuth is selected, and synapses from active KCs (*i*) onto this MBON (*j*) are depressed according to the following anti-hebbian rule:

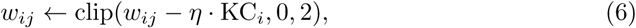

with learning rate *η* = 0.5. This ensures that each snapshot is associated with the appropriate MBON. During inference, all MBONs are evaluated in parallel, and the MBON with the lowest activation—corresponding to the most familiar directional prediction—is selected. The egocentric azimuth associated with the selected MBON defines the predicted nest azimuth.

KC activity patterns were first stored in a short-term buffer and the KCs to MBON synapses are depressed (consolidated) into long-term memory after return to the nest. The precise mechanism of visual memory consolidation may vary, but a key requirement of our model is that newly acquired snapshots are not immediately used for navigation, as this would disrupt spiralling behaviour.

### Error-driven control of LWs

The instantaneous PI–Vision disagreement (*E*_PI*/*V_) is computed as:

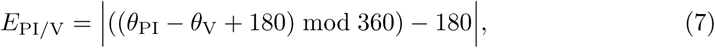

and normalized between 0 and 1 through a cosine kernel:

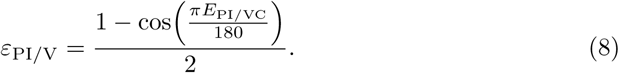

**Table 1.**
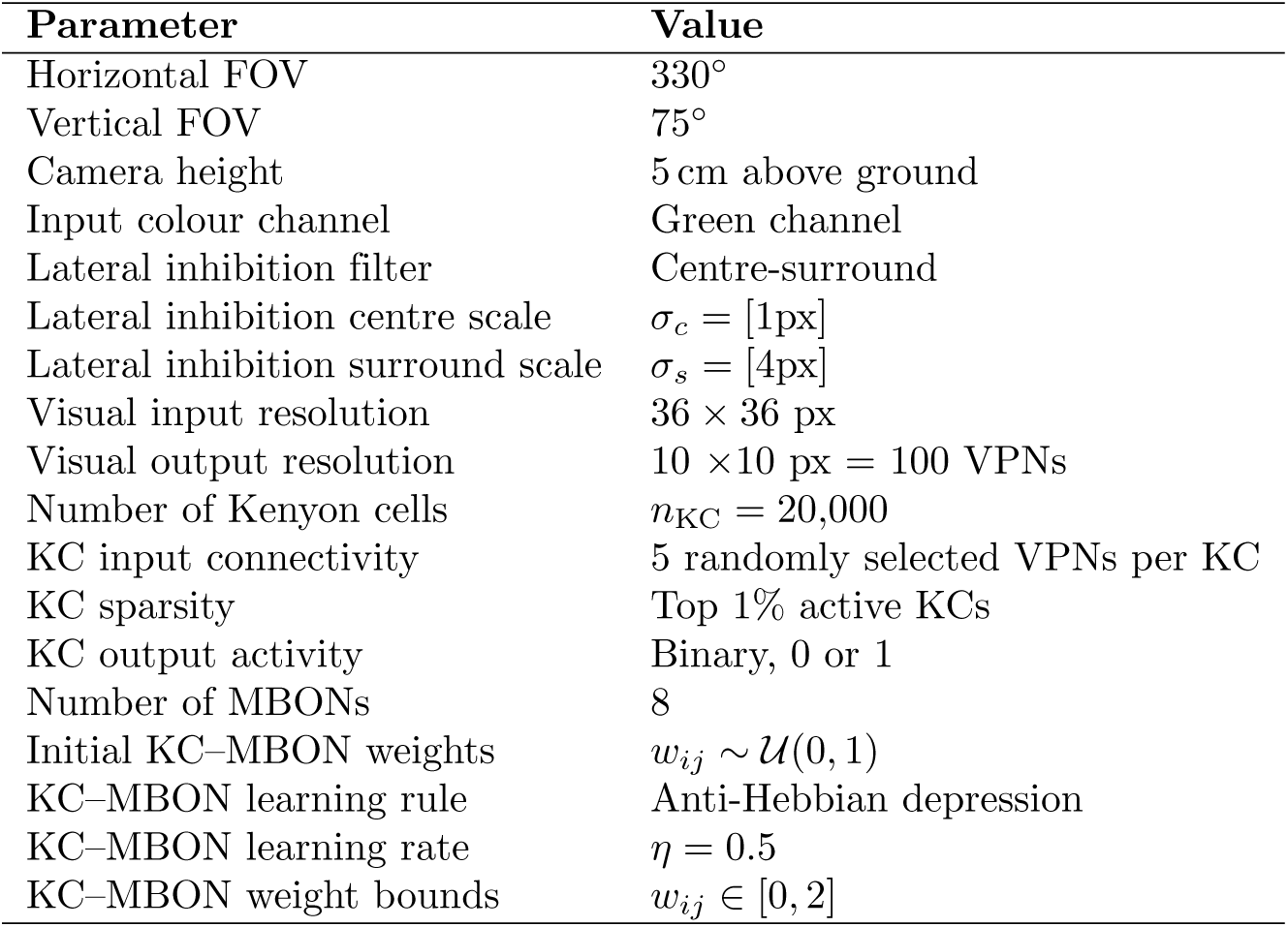
Vision preprocessing and mushroom body visual compass parameters used in the model.

As learning progresses, it is expected that *E*_PI_*_/_*_VC_ decreases in previously explored areas. The cumulative PI–Vision error *C*_PI_*_/_*_VC_ is defined as the running sum of instantaneous angular errors across a single LW:

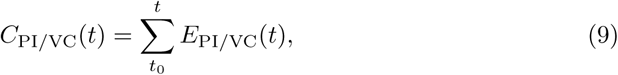

which is reset after coming back to the nest. The instantaneous PI–Vision disagreement *ε*_PI_*_/_*_VC_ ∈ [0, 1] modulates navigation and learning through the following mechanisms:

### Steering and spiraling

At each timestep, the agent computes a goal direction based on its current navigational mode (*inward* or *outward* ). For inward navigation, the agent steers toward the predicted nest azimuth (*θ*_PI_); for outward navigation, the goal direction is set to (*θ*_PI_ + 180*^◦^*). At the onset of walks, the agent follows the outward direction.

The nest azimuth is estimated from a combination of PI and visual compass (VC), yielding a single heading *θ*_VCPI_. This fusion is performed via circular vector averaging:

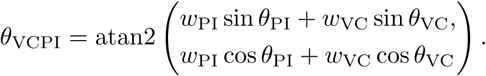

Here, *w*_PI_ = 0.4 and *w*_VC_ = 0.6 in all experiments. In real ants, this combining of visual and PI cues, probably happens in CPU1 neurons of the CX. These weights were chosen so that PI dominates during early learning, when the visual compass has a low signal-to-noise ratio, while still allowing the visual compass to overcome the PI signal once its predictions become more accurate and consistent.

The agent moves forward at each timestep and cannot move backward or sideways; steering is achieved by rotating its heading toward the current goal direction, with a maximum angular change of 30*^◦^* per timestep. The turning angle is computed as the signed circular difference between the agent’s current heading and the goal direction, clipped to this maximum turn rate and corrupted by zero-mean Gaussian noise with standard deviation *σ* = 15*^◦^*. LWs have been described as arcs around the nest, with the ant walking perpendicular to the nest [54]. To promote spiraling behaviour during early LW, the agent momentarily switch goal direction and follow a goal perpendicular to the homing vector. This occurs with probability

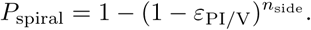

At the onset of LWs, a random spiraling direction is selected (clockwise or counterclockwise relative to the vertical axis, *p* = 0.5). The parameter *n*_side_ determines the likelihood of triggering spiraling motion and is set to ten in all experiments

### Pirouette and snapshot acquisition

At each timestep of the LWs, a pirouette is triggered with probability

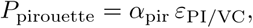

where *α*_pir_ is set to 0.15 in all experiments. At the onset of each pirouette, the agent is randomly assigned to perform either a *partial* or *full* pirouette, with *p* = 0.1 for full pirouettes. The agent rotates in discrete 1*^◦^* steps in a randomly chosen direction (*p* = 0.5). Partial pirouettes are terminated when the heading reached the nest direction within ±5*^◦^*; full pirouettes lasted 360*^◦^*.

During rotation, an accumulator is updated at each step with a random value sampled from *Unif* [0, 1]. At each step, the snapshot threshold is drawn from N (*µ* = 65*, σ* = 35), with non-positive values rejected. When the accumulator exceeds this threshold, a snapshot is acquired and the accumulator is reset. Snapshots therefore occurred stochastically and independently of absolute angular position.

We optimised parameters so that pirouettes contain on average 4 ± 3 fixations, matching observations in *Cataglyphis noda* [59] (Fig. 13B). After each pirouette, the agent’s body orientation is reset to its pre-pirouette heading, so pirouettes affect memory acquisition but not the walk trajectory. Successive fixations were separated by approximately 45*^◦^* on average (Fig. 13A), supporting uniform sampling of the surrounding landscape (Fig. 13E,F).

### Premature return

The cumulative disagreement between PI and visual cues is updated at each timestep as

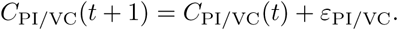

When the cumulative error exceeds a fixed threshold of 125, the agent switches from outward to inward navigation mode, triggering return to the nest.

### Walking speed modulation

After turning, the agent moves forward by a distance *v*, defined as *v* = 1 − *E*_PI_*_/_*_VC_ + 0.1 cm/step. This results in slower walking speed with higher *E*_PI_*_/_*_VC_. There is a residual speed of 0.1 centimeter per timestep to prevent the agent becoming stuck when the error is 0. The maximum speed is effectively 1.1 centimeter per timestep. In real ants, slower walking speed may support increased sampling and evidence accumulation. In our model, however, speed modulation plays no explicit functional role beyond reproducing the experimental observation that LWs are associated with lower walking speeds than foraging walks.

### Food picking

Food acceptance is implemented as a thresholded response based on the recent PI and visual cues mismatch, *E*_PI_*_/_*_VC_, averaged over the last 100 timesteps. At the release point, located three meters from the nest, food is accepted if the recent average of *E*_PI_*_/_*_VC_ falls below a fixed threshold. Lower PI-VC mismatch promotes food acceptance.

**Table 2.**
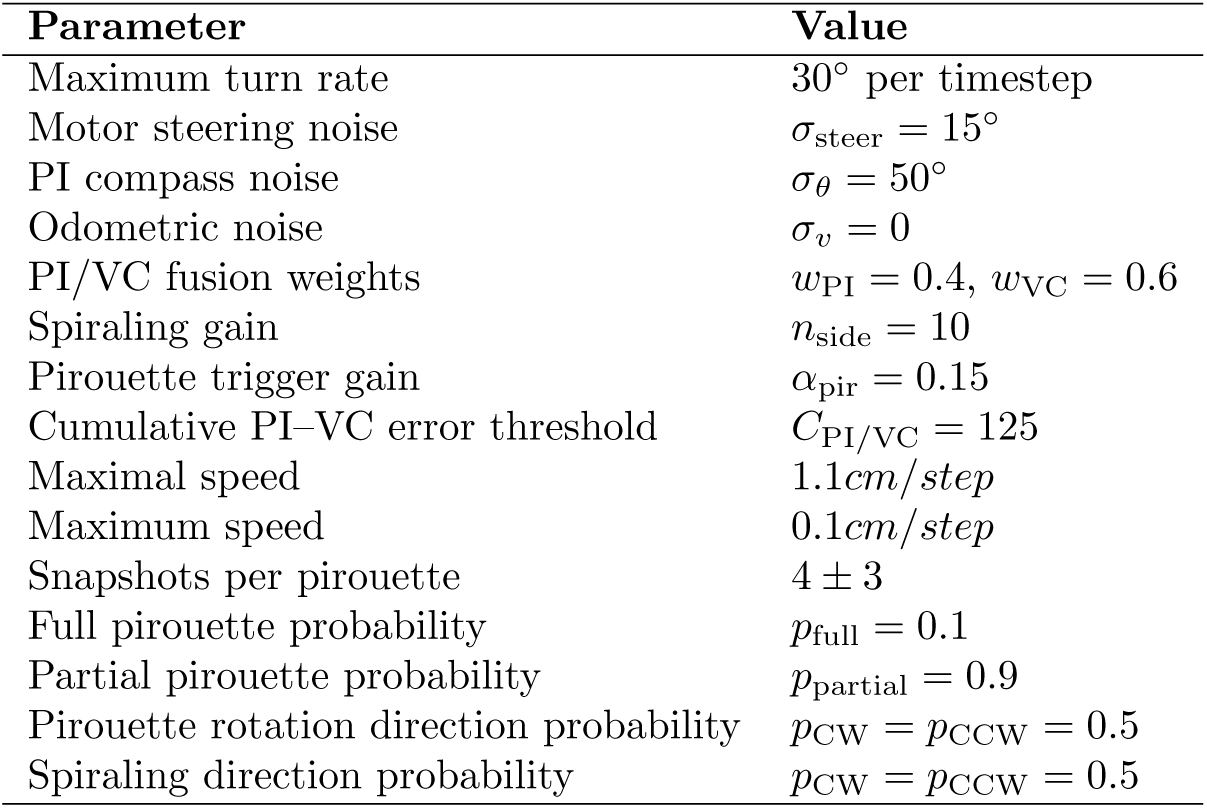
Parameters were hand-tuned to replicate experimental behaviours across all three experimental layouts.

## 1 Supplementary Information

### 1.1 Clawed walks / Relearning walks

#### Introduction and results

Jayatilaka et al. [58] reported that even experienced foragers occasionally perform short, LW–like segments near the nest, before starting their foraging walks (Fig. 11A). They termed these ‘clawed walks’ and this behaviour is also present in our model (example trajectories are shown in Fig. 11B). We find that 38% of the foraging walks (maximum distance to nest *>* 3 m) form these claws in our model. For both PI and visual prediction, angular error relative to the true nest position was significantly higher in the 0–10 cm distance bin compared to all other bins (Welch’s *t*-tests with FDR correction, all *p <* 0.001; Fig. 11C). This analysis included only trajectories where agents had ventured beyond 5 m from the nest, ensuring the comparison was restricted to experienced agents.

**Fig 11.**
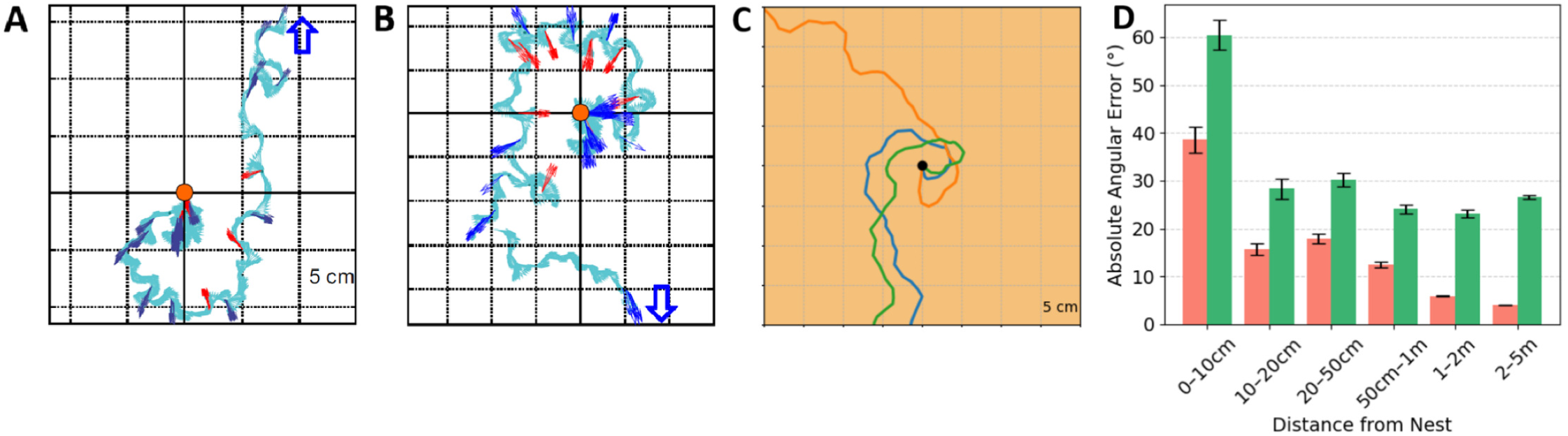
Clawed walks. **A,B**: Empirical examples from [58], showing foraging walks that begin with a clawed loop around the nest. **C,D**: The same behaviour emerges in the model, occurring in approximately 38% of foraging walks. **E**: Angular error between PI predictions and the true nest direction (red bars) across six distance bins; the 0–10 cm bin shows markedly higher error than all others. **F**: Angular error between visual predictions and the true nest direction (green bars); even after training, errors remain elevated at very close range to the nest.

#### Discussion

Jayatilaka et al. [58] interpreted clawed walks as partial learning segments that may provide contrastive views opposite to the future foraging direction, reducing the risk of entering unfamiliar terrain after overshooting the nest. In our model, clawed walks emerge without an explicit functional purpose. They arise as a byproduct of the spiralling rule combined with elevated PI–visual error in the immediate nest vicinity (0–10 cm), where both PI and visual predictions are substantially less reliable than at larger distances (Fig. 11).

### 1.2 Prediction of the model: Displacement-induced PI–vision conflict

#### Introduction and Results

This experiment tests a prediction generated by our model that has not yet been verified experimentally. If the mechanisms implemented in the model reflect those operating in the insect brain, ants should display similar behaviour. PI is sensitive to displacement, whereas visual predictions remain anchored to the surrounding panorama. Therefore, grabbing an experienced forager near the nest and releasing it on the opposite side generates a mismatch between PI- and vision-based estimates of nest direction that increases with angular offset (Fig. 12A).

**Fig 12.**
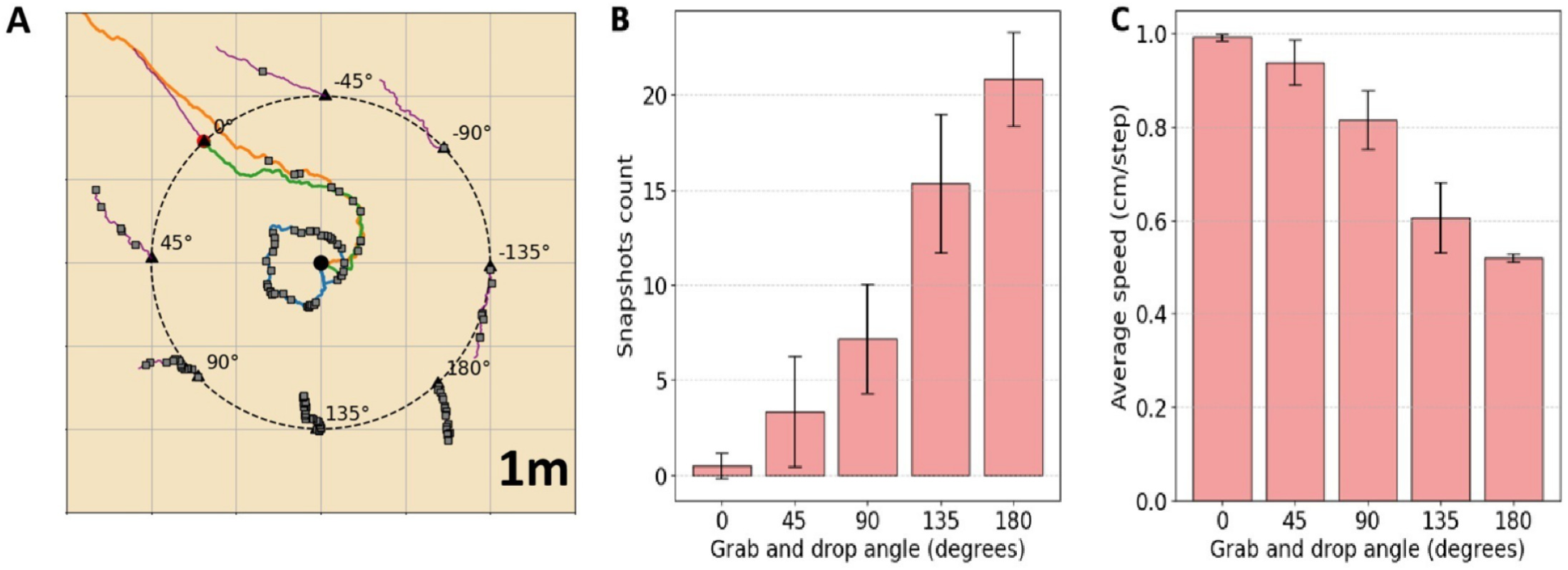
Prediction experiment: Displacement-induced PI–vision conflict **A:** Example of one complete training and test protocol. Nest at the center. Grabbing point in red (0°). Blue line: first LW; orange line: second LW. Grey squares: pirouettes. Green line: third walk and pre-grab trajectory, stopping at the grabbing point, 2 m from the nest. Purple lines: post-drop walks starting at different locations 2 m around the nest. **B:** Number of snapshots recorded during the post-drop walks. Larger angular displacement results in a higher number of pirouettes. **C:** Speed decreases with increasing angular displacement.

**Fig 13.**
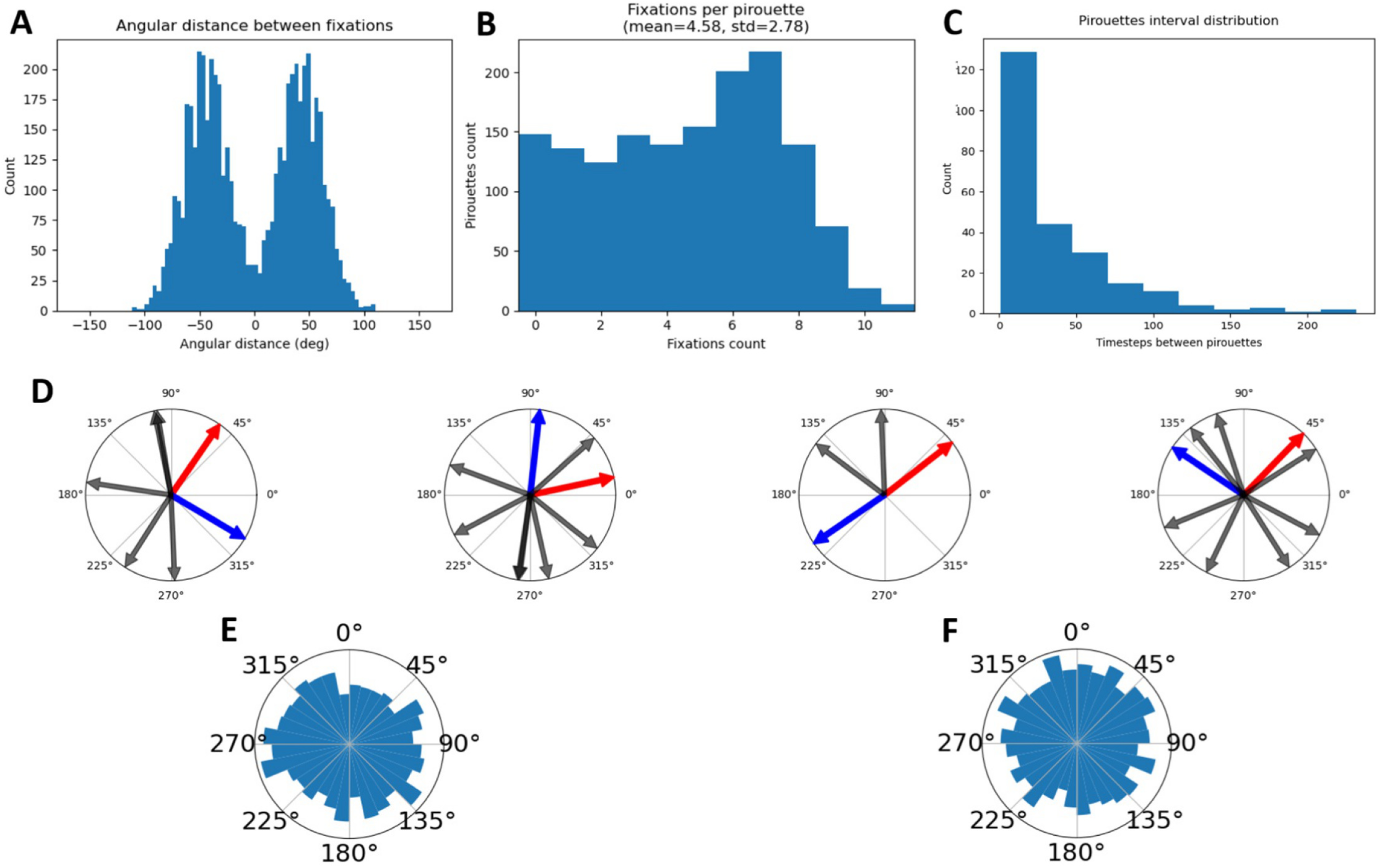
**A:** Angular interval between fixations (snapshots). The mean angular distance between fixation is 45 degrees. (During replication of Müller and Wehner experiment) **B:** Distribution of the number of snapshots per pirouette. Mean = 4.58 and Standard deviation = 2.78. **C:** Exponentially delayed interval between pirouettes. **D:** Four examples of pirouettes. Blue arrows: initial agent azimuth. Red arrows: nest azimuth. Black arrows: Snapshots directions **E:** Nest-centric distribution of the snapshots. Facing nest at the top (0deg). **F:** Allocentric distribution of the snapshots. Facing north at the top (0deg). In both frame of reference, the distribution is broad and uniform and does not show a bias to a particular direction.

Our model continuously computes this PI–vision mismatch in real time and therefore predicts that such displacements will reactivate LW–like behaviour. To test this prediction, agents were first trained through successive LWs until they reached the foraging phase. Foraging agents were grabbed at a distance of 2 m from the nest and released at different locations on that 2 m radius ring, which introduced controlled angular error between the PI-predicted nest direction and the true nest direction (0°, ±45°, ±90°, ±135°, 180°), without affecting the visual-compass prediction.

The number of snapshots recorded during post-drop walks increases monotonically with angular displacement (Fig. 12B; Spearman *ρ* = 0.911, *p* = 2.69 × 10*^−^*^12^), while both walking speed (Fig. 12C; *ρ* = −0.926, *p* = 2.34 × 10*^−^*^13^) and path straightness (Fig. 12D; *ρ* = −0.939, *p* = 1.49 × 10*^−^*^14^) decrease monotonically.

#### Discussion

The model predicts that displacing an experienced forager after it has left the nest, while preserving its visual input but rotating its PI estimate, should reactivate LW–like behaviour. The magnitude of this response is predicted to scale with the angular mismatch introduced between the PI-based and visually based estimates of nest direction. Wystrach et al. [69] already showed in *Melophorus bagoti* that scanning reoccur in experienced foragers when PI conflicts with visual cues but, extending this experiment as we suggest would provide direct behavioural evidence as to whether the mismatch between PI and vision controls LW dynamics in the way we hypothesise.

